# M. tuberculosis invades and disrupts the blood brain barrier directly to initiate meningitis

**DOI:** 10.1101/2024.08.30.610462

**Authors:** Alizé Proust, Katalin A Wilkinson, Robert J Wilkinson

## Abstract

Tuberculous meningitis (TBM) is the most severe form of tuberculosis and HIV-1 co-infection worsens the already poor prognosis. How *Mycobacterium tuberculosis* (*Mtb*) crosses the blood brain barrier (BBB) and the influence of HIV-1 on pathogenesis remains unclear. Using human pericytes, astrocytes, endothelial cells, and microglia alone; and combined in an *in vitro* BBB we investigated *Mtb* +/− HIV-1 co-infection on central nervous system cell entry and function. *Mtb* infected and multiplied in all cell types with HIV-1 increasing entry to astrocytes and pericytes, and growth in HIV-1 positive pericytes and endothelial cells. The permeability of the BBB increased resulting in translocation of bacilli across it. Cytopathic effects included increased markers of cellular stress, ROS release, the induction of neurotoxic astrocytes, and increased secretion of neuroexcitotoxic glutamate. Distinct cell-type specific production of inflammatory and effector mediators were observed. These data indicate *Mtb* can translocate the BBB directly to initiate meningitis.

## Text

Tuberculosis (TB), caused by *Mycobacterium tuberculosis* (*Mtb*) is a major global cause of morbidity and mortality, affecting millions of individuals each year^1^. TB is significantly exacerbated by HIV-1 co-infection and this combination is especially important in regions with a high prevalence of both diseases^2^. *Mtb* enters via the lungs (pulmonary TB) but can also disseminate throughout the body (extrapulmonary TB), including to the central nervous system (CNS), leading to meningitis (TBM). TBM accounts for approximatively 1-10% of all active TB cases, depending of the local TB prevalence, and is the most destructive manifestation of extrapulmonary TB, resulting in severe morbidity and high mortality^3–6^.

The mechanisms by which *Mtb* transmigrates to the brain are not fully understood: both lymphatic and hematogenous spread have been suggested^3^. Once in the systemic circulation, *Mtb* could enter the brain as cell-free bacilli across the blood-brain barrier (BBB) or blood-cerebrospinal-fluid barrier (BCSFB), be trafficked across the BBB in infected neutrophils and macrophages^7^, or disrupt endothelial tight junctions to enter between cells. The BBB, principally composed of brain microvascular endothelial cells forming tight junctions, prevents most pathogens from entering the brain. *Mtb* has however been shown to invade the brain through rearrangement of actin of BBB endothelial cells, with *Mtb* pknD gene necessary to invade these cells^8, 9^.

HIV-1 is a risk factor for extrapulmonary TB and TBM with co-infection associates with a poor prognosis^10^. While the clinical association between HIV-1 and TB is well-characterized^1^, the cellular effects of co-infection of brain cells remain poorly understood. This study evaluated the entry and effect of *Mtb* on CNS cells and the BBB and characterised the effect of co-infection with HIV-1 on this process. We infected astrocytes, pericytes, brain endothelial cells and microglia and an *in vitro* BBB model with HIV-1 and/or *Mtb* and analysed their effects on CNS cells and BBB function. We showed that *Mtb* effectively enters and replicates in CNS cells causing cytotoxicity and loss of integrity of the BBB. *Mtb* induced reactive oxygen species release, endoplasmic reticulum stress and astrogliosis; affected mitochondrial activity; increased extracellular glutamate concentration and led to pro-inflammatory cytokine and chemokine secretion. Although Mtb tended to dominate effects, prior infection with HIV-1 aggravated its effects depending upon cell type. Our results throw light on the impact of *Mtb* and HIV-1 on CNS cells and BBB function and its potential amelioration by novel antibiotic and anti-inflammatory treatment strategies.

## Results

### Effect of HIV-1 infection on *Mtb* entry and growth in CNS cells

Apart from microglia/macrophages, brain cells are poorly susceptible to HIV-1 infection and the microglia cell line (HMC3) used does not express the CD4 receptor. We therefore used both HIV-1_BaL_ and a VSV-G pseudotyped HIV-1 virus in our study. HIV-1_BaL_ allowed us to study bystander effects of HIV-1 on CNS cells while VSV-G pseudotyped HIV-1 was used to determine the effect of productive infection with HIV-1.

We first investigated whether HIV-1 infection modulated *Mtb* uptake into CNS cells. Cells were incubated with HIV-1 (BaL or VSV-G pseudotyped) for 24 hrs before *Mtb* infection (3 hrs), with washing to removed unbound bacteria, and staining. Flow cytometric analysis of fluorescent bacteria was used to assess uptake and growth, and the gating strategy is shown in Supplementary Figure 1. HIV-1 pre-infection facilitated *Mtb* entry in astrocytes (Fig. 1A, *Mtb*+: 7.5 ± 1.2%; BaL + *Mtb*: 8.9 ± 1.4%; VSV-G + *Mtb*: 9.4 ± 1.7%), pericytes (Fig. 1E, *Mtb*+: 7.5 ± 1.0%; BaL + *Mtb*: 14.9 ± 1.1%; VSV-G + *Mtb*: 15.4 ± 1.2%) and microglia (Fig. 1G, *Mtb*+: 7.1 ± 1.2%; BaL + *Mtb*: 7.7 ± 1.3%; VSV-G + *Mtb*: 8.8 ± 1.6%). Endothelial cells were also infected by *Mtb* but HIV-1 infection had no effect on their susceptibility to bacteria (Fig. 1C, *Mtb*+: 17.1 ± 3.6%; BaL + *Mtb*: 16.7 ± 3.4%; VSV-G + *Mtb*: 17.5 ± 3.5%). No effect, however, on the MFI of *Mtb*+ astrocytes, pericytes, microglia and endothelial cells between *Mtb*-infected cells and *Mtb*/HIV co-infected cells was observed (Fig. 1B,D,F,H).

**Figure 1.**
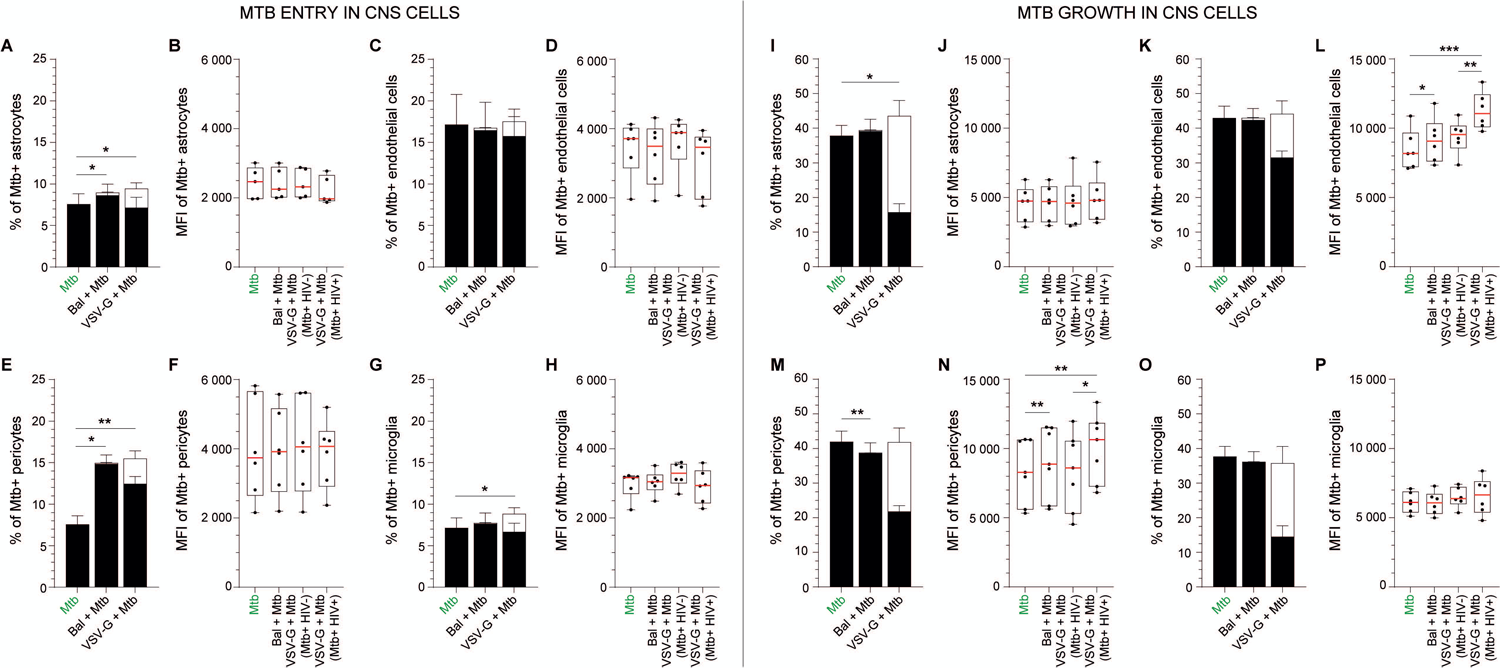
*Mtb* entry and growth in CNS cells. (A, B, I, J) Astrocytes, (C, D, K, L) hCMEC/d3 (endothelial cells), (E, F, M, N) HBVP (pericytes) and (G, H, O, P) HMC3 (microglia) were either left uninfected, infected with HIV-1 Bal or VSV-G pseudotyped HIV-1 for 24 hrs before being incubated for 3 hrs with *Mtb* followed by extensive washing. Cells were then directly stained and analyzed by flow cytometry (*Mtb* entry in CNS cells, A-H) or incubated for 48 hrs. (*Mtb* growth in CNS cells, I-P). (A, C, E, G, I, K, M, O) The percentage of *Mtb*+ and *Mtb*+HIV+ cells are displayed in filled and opened bars, respectively. (B, D, F, H, J, L, N, P) The MFI of *Mtb*+ cells is displayed for both infection conditions (*Mtb*+ and *Mtb*+HIV+). The mean and SEM (A-H) and medians (I-P) for 5 to 6 different donors/passages are shown. Asterisks denote statistically significant data as determined by one-way analysis of variance (ANOVA) with corrections for multiple comparisons (Tukey) (\**P* < 0.05, \*\**P* < 0.01, \*\*\**P* < 0.001)

Interestingly, *Mtb* appeared to preferentially infect HIV negative cells (Fig. 1A,C,E,G: filled bars versus opened bars). Only 23.0 ± 4.4% of *Mtb* positive astrocytes were HIV-1 co-infected (Fig. 1A). This percentage dropped to 20.4 ± 5.1% in microglia (Fig. 1G), 18.5 ± 4.2% in pericytes (Fig. 1E) and 9.3 ± 2.3% in endothelial cells (Fig. 1C).

To assess *Mtb* growth in CNS cells, we incubated cells for 3 hrs followed by washing off cell-free bacilli and a further 48 hrs incubation. The percentage of *Mtb*+ cells increased after 48 hrs (*Mtb*+ astrocytes 37.8 ± 3.0%; pericytes 41.9 ± 3.1%; endothelial cells 42.9 ± 3.4 %; and microglia 37.6 ± 3.0%, Fig. 1I-P). Growth was slightly but significantly increased in astrocytes by productive HIV-1 infection (VSV-G + *Mtb*: 43.5 ± 4.1 %, Fig. 1I). By contrast the MFI of *Mtb*+ astrocytes and microglia remained unchanged between *Mtb*-infected and *Mtb*/HIV-co-infected cells (Fig. 1J and P). The MFI of *Mtb*+ pericytes (median MFI: 8281, IQR 5591–10632) and endothelial cells (MFI: 8173, IQR 7207 – 9668) was significantly increased in *Mtb*+ cells in contact with HIV-1_BaL_ (pericytes: 8865, IQR 5844–11497, endothelial cells: 9073, IQR 7611 – 10342, Fig. 1L,N) and *Mtb*+ cells productively infected with VSV-G pseudotyped HIV-1 (pericytes: 10639, IQR 7257– 11837; endothelial cells: 11055, IQR 10089–12440, Fig. 1L,N).

### Effect of HIV-1 infection on the *Mtb*-induced BBB permeability and *Mtb* translocation

We then investigated the impact of both infections on a 3D human BBB model, in which permeability was assessed by measuring the diffusion of dextran-rhodamine B from the upper chamber to the collector after two or six days post-*Mtb* infection.

Infection with *Mtb* increased BBB permeability after two (and more so after six) days of infection. The additional impact of HIV-1 infection on BBB integrity was however negligible (Fig. 2A,B). Consistent with the breakdown of the BBB, *Mtb* could be recovered in the collector after 2 days of infection (median 20 CFU/mL) which was approximately doubled by *Mtb*/HIV-1_Bal_ (47 CFU/mL, Fig. 2C). After six days’ culture the concentration of *Mtb* in the collector increased greatly (median 7359 CFU/mL), although values were not further increased by either form of HIV-1 co-infection (Fig. 2D).

**Figure 2.**
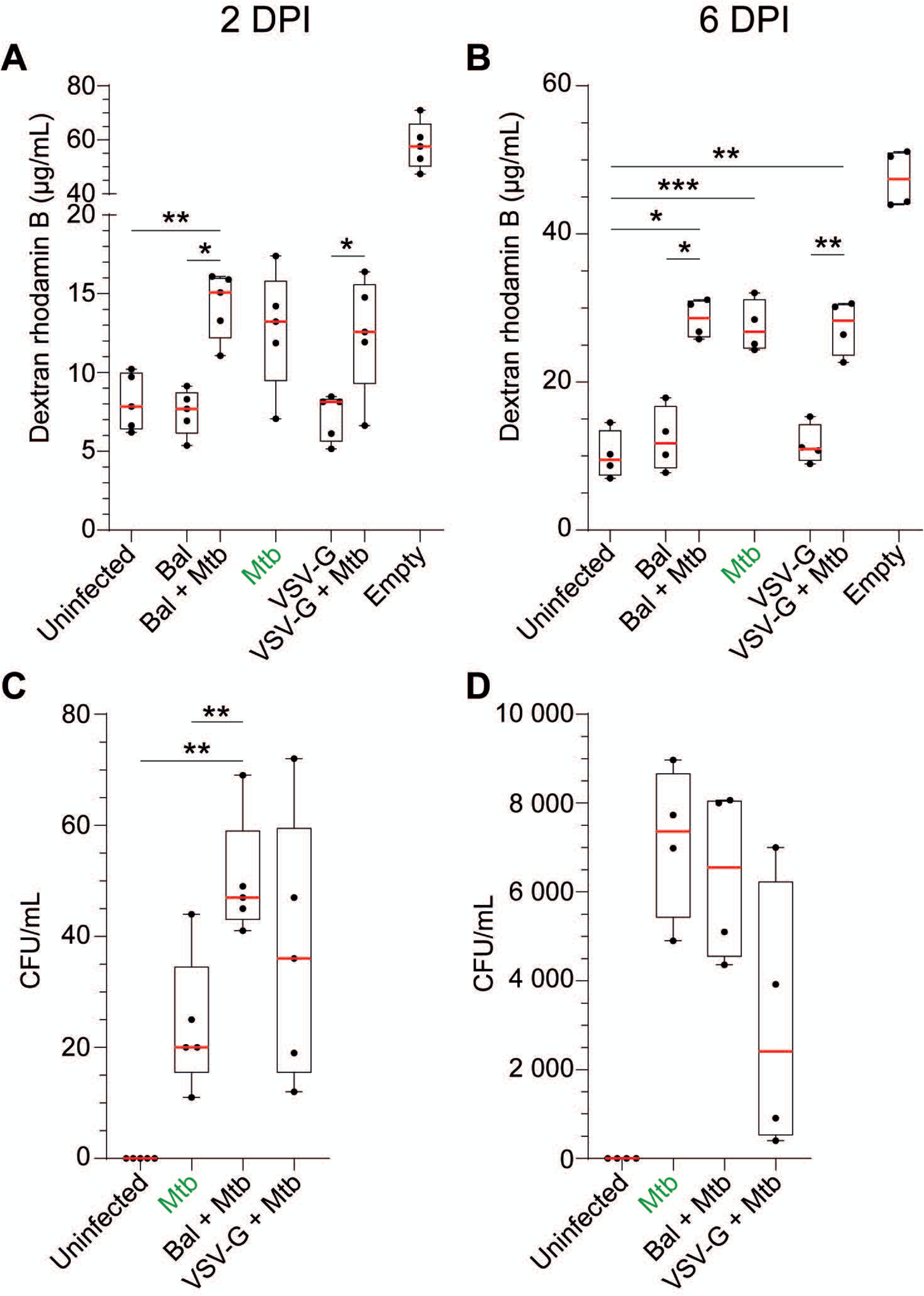
BBB permeability and *Mtb* passage through the BBB. BBB were infected with HIV-1 for 24 hrs. and/or *Mtb* for (A, C) 2 days or (B, D) 6 days and we measured (A, B) BBB permeability (dextran-rhodamine assay) and (C, D) *Mtb* concentration in the collector (CFU). An empty control insert (“empty”) was used to estimate the passive diffusion of dextran-rhodamine from the upper chamber to the collector and mimicked a BBB with 100% permeability. Data of 4 to 5 experiments are presented in raw data (i.e., concentration of Dextran-rhodamine B 70kDa in µg/mL or concentration of *Mtb* in CFU/mL) in box and whiskers plots. Asterisks denote statistically significant data defined by ANOVA with Tukey’s correction for multiple comparisons (*P<0.05, **P<0.01).

### Effect of *Mtb* and HIV-1 infections on CNS cell monolayer integrity and tight junction expression

We next assayed in real time the effect of *Mtb* and HIV-1 infections on astrocyte, pericyte, and microglial monolayers as well as on brain microvascular endothelial barrier integrity using the impedance-based xCELLigence RTCA system (Fig. 3). During 80 hrs *Mtb* steadily decreased the impedance of all cell types by over 80%. Productive HIV-1 infection alone decreased the impedance of astrocytes and pericytes. Combined infection effects were not greater than *Mtb* alone except in productively HIV-1 and *Mtb* infected pericytes (Fig. 3F).

**Figure 3.**
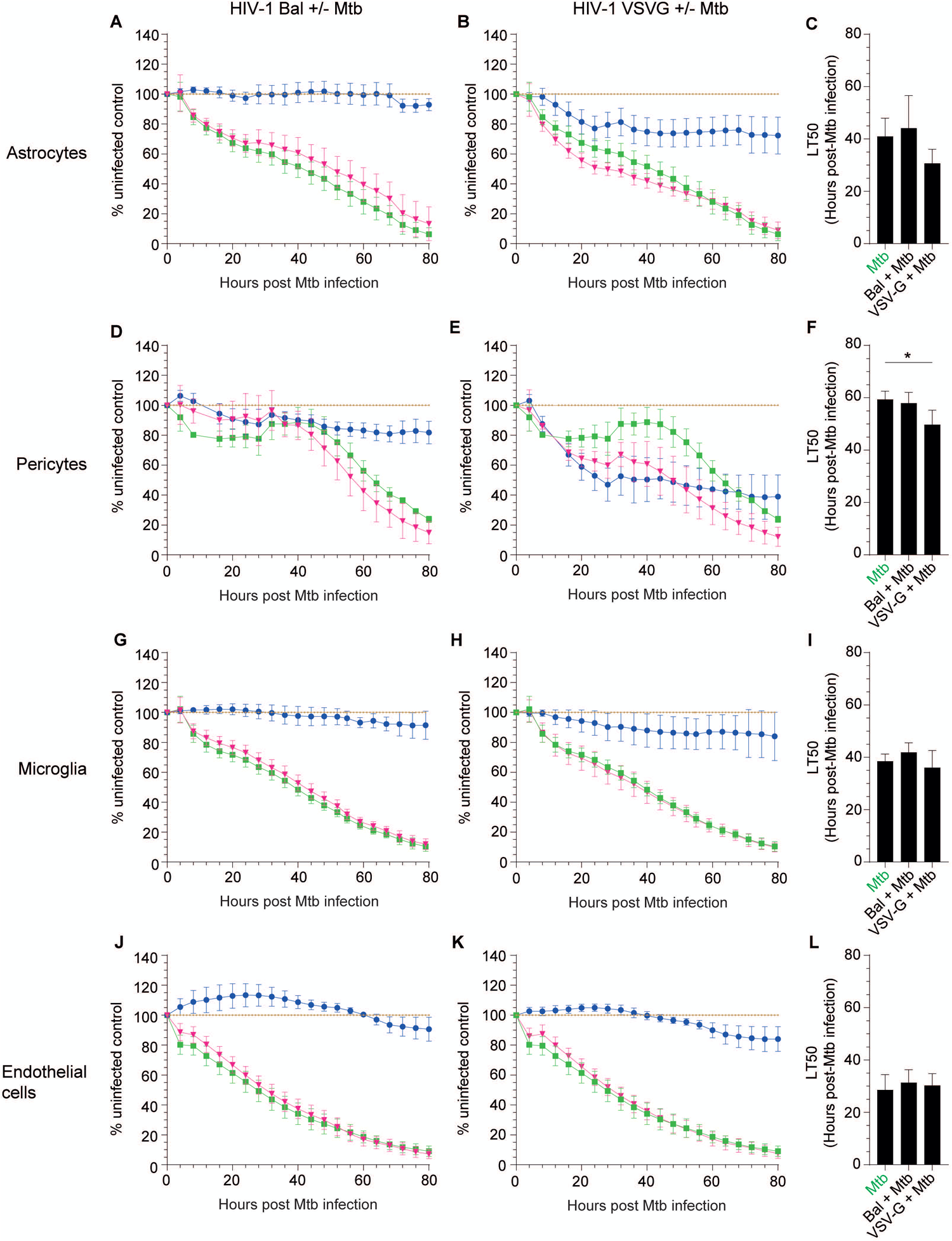
*Mtb* and HIV-1 cytopathic effect on CNS cells. Astrocytes (panels A-C), HBVP (D-F), HMC3 (G-I) and hCMEC/d3 (J-L) were either *Mtb*-mono-infected (green, A, B, D, E, G, H, J, K), HIV-1 Bal mono-infected (blue, A, D, G, J), VSV-G pseudotyped HIV-1 mono-infected (blue, B, E, H, K), *Mtb*/HIV-1 Bal co-infected (pink, A, D, G, J) or *Mtb*/VSV-G pseudotyped HIV-1 co-infected (pink, B, E, H, K) and the impedance was recorded in real time continuously for 3 days post-*Mtb* infection. The mean and SEM for 4 to 5 different donors/passages are presented compared to uninfected controls (brown). (C, F, I, L) The lethal time 50 (time at which impedance is reduced 50%) for 4 to 5 different donors/passages is shown as mean and SEM. Asterisks denote statistically significant data as determined by ANOVA with Tukey’s correction for multiple comparisons (\**P* < 0.05).

We also assayed endothelial cell expression of transcripts for tight junction proteins (ZO-1, JAM-A, Occludin and Claudin-5) and adherens junction protein (VE-Cadherin) in response to HIV-1, *Mtb* and *Mtb*/HIV-1 infections (Supplementary Figure 2). Overall, effects were moderate and no statistically significant changes were detected in the presence of HIV-1, *Mtb* or during co-infection.

### Effect of *Mtb* +/− HIV infection on mitochondrial metabolic activity and reactive oxygen species (ROS) release

We next measured the effect of both infections on CNS cell mitochondrial metabolic activity and ROS release. Apart from a small but significant decrease in mitochondrial activity of endothelial cells conferred by *Mtb* and productive HIV-1 co-infection (1.16-fold median decrease *by Mtb* and slightly worsened by co-infection to 1.30-fold, Fig. 4B), there was no significant effect of either infection on other cells (Fig. 4A,C,D).

**Figure 4.**
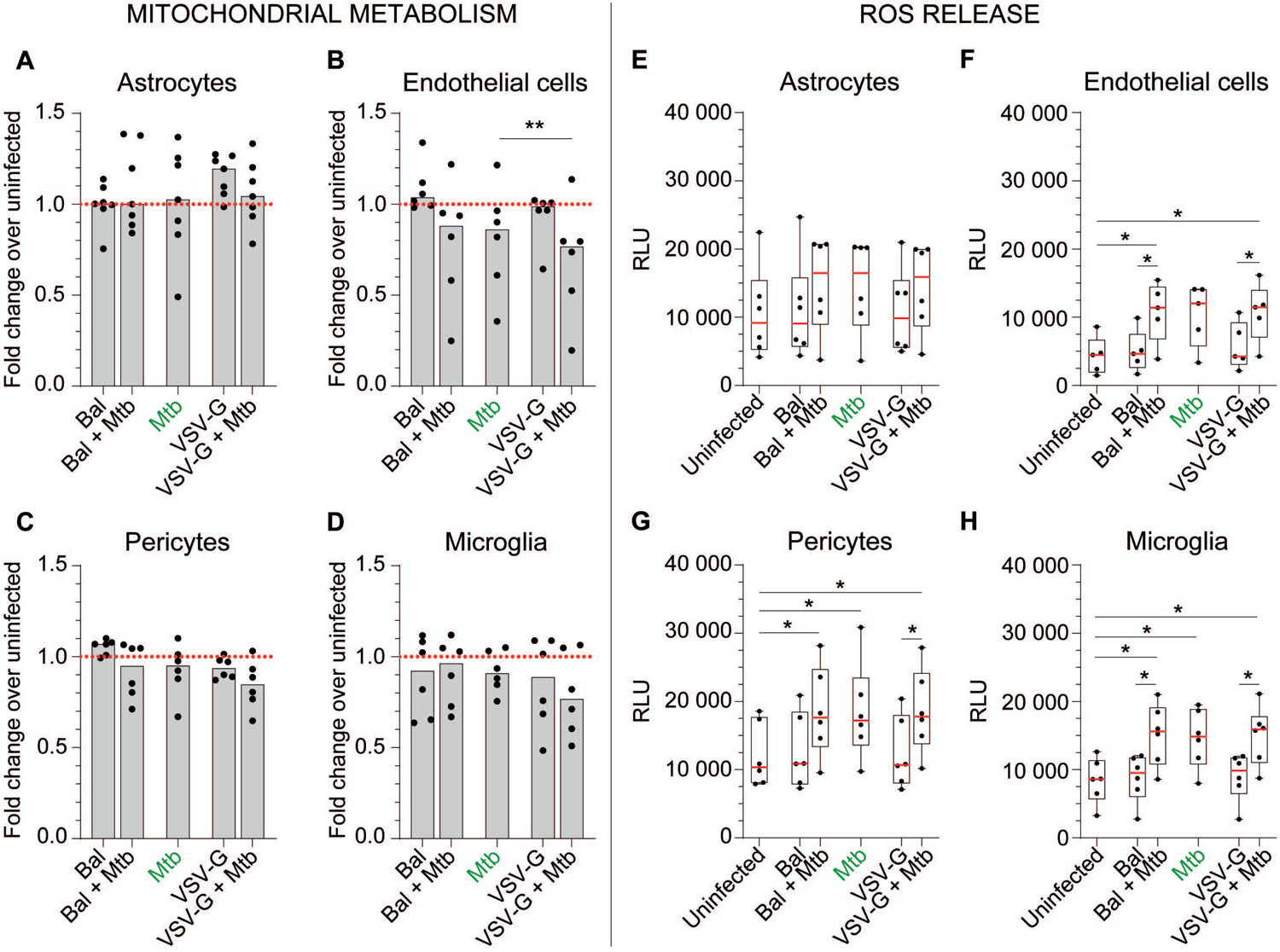
Effect of *Mtb* and/or HIV-1 infection(s) on CNS cell metabolic activity and ROS release. Mitochondrial metabolic activity (MTS assay, panels A-D) and the release of H_2_O_2_ (ROS assay, panels E-H)) by CNS cells were assessed for astrocytes (A,E), hCMEC/D3 (B,F), HBVP (C,G) and HMC3 (D,H) following infection(s) with *Mtb*, HIV-1 Bal and/or VSV-G pseudotyped HIV-1. The medians for 5 to 6 different donors/passages are presented in fold change over the uninfected condition (dashed red line, MTS assay) or in raw data in box and whiskers plots (ROS assay). Asterisks denote statistically significant data as determined by ANOVA with Tukey’s correction for multiple comparisons (\**P* < 0.05, \*\**P* < 0.01)

When treated with 50 µM menadione as a positive control, all cell types increased ROS release more than ten-fold (Supplementary Figure 3). *Mtb* also increased ROS release by all CNS cell types up to two-fold, significantly so for pericytes and microglia (Fig. 5G,H). Neither bystander or productive HIV-1 infection further increased *Mtb*-induced ROS release (Fig. 2E-H).

**Figure 5.**
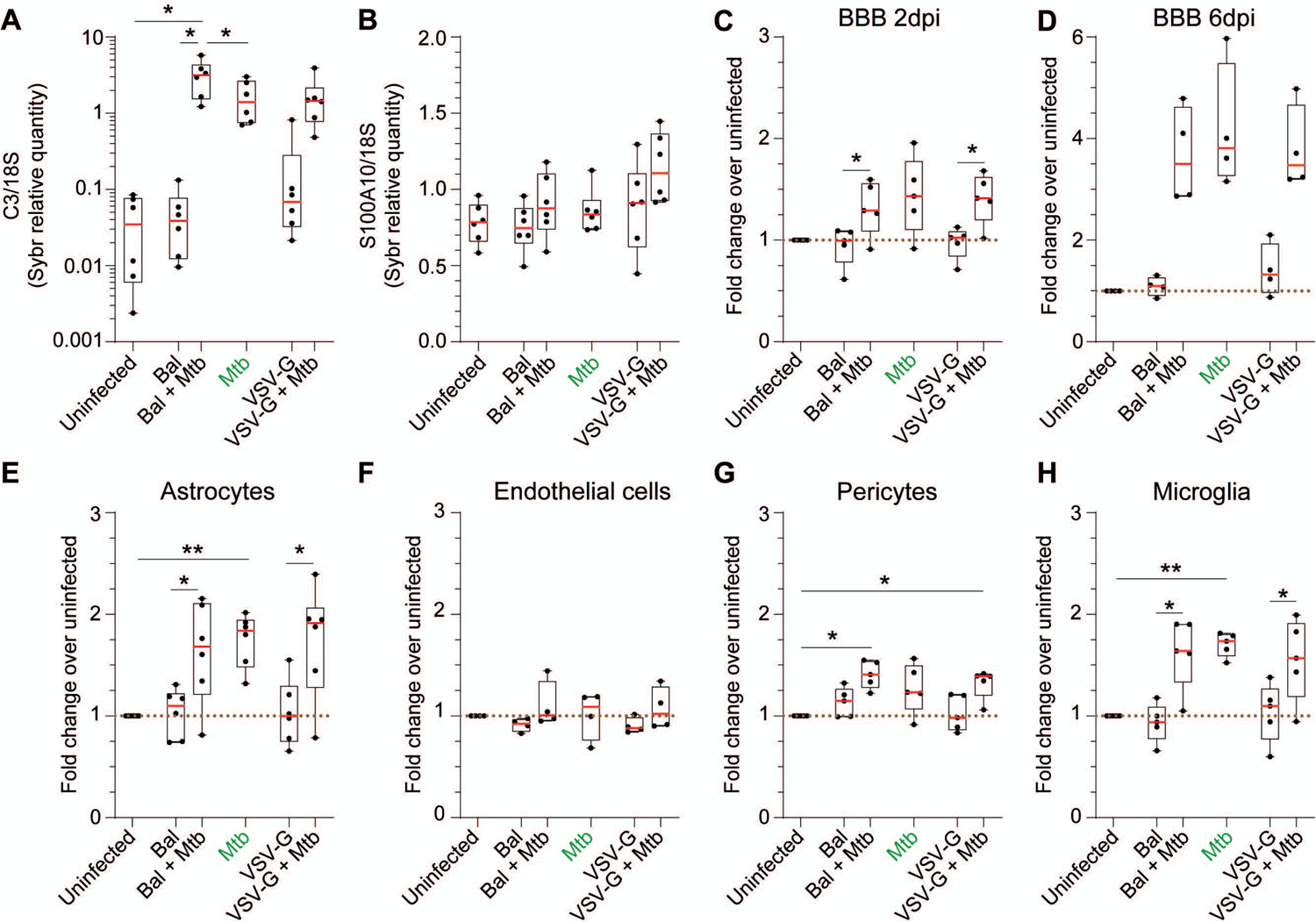
Effect of *Mtb* and/or HIV-1 infection(s) on astrogliosis and extracellular glutamate concentration. (A) Complement 3 (C3) and (B) S100A10 gene expression was assessed by SYBR-Green quantitative RT-PCR following infection(s) with *Mtb*, HIV-1 Bal and/or VSV-G pseudotyped HIV-1. Results for 5 different donors are presented as raw data normalized to the geometric mean of 18S ribosomal gene expression. CNS cells and BBB were infected as previously described for 2 (C, E, F,G,H) or 6 days (D) and extracellular glutamate was determined. Data of 4 to 5 experiments are presented as fold change over the uninfected control (brown dashed line). Asterisks denote statistically significant data analysed by ANOVA with Tukey’s correction for multiple comparisons (*P<0.05, **P<0.01).

### Effect of *Mtb* infection on the unfolded protein response (UPR)

To expand investigation of *Mtb* and HIV-1 infections on cellular stress, we next investigated the effect on endoplasmic reticulum stress by evaluation of transcript levels within three UPR signalling pathways (Supplementary Figure 4). The splicing of XBP1 was increased in all CNS cell types following *Mtb* infection (median increase in astrocytes: 1.35-fold; endothelial cells: 7.62-fold; pericytes: 2.63-fold; microglia: 3.75-fold, Supplementary Figure 4A-D). Productive HIV-1 infection significantly augmented this increase in microglia (5.45-fold) and bystander co-infection in pericytes (3.21-fold, Supplementary Figure 4C,D). By contrast, *Mtb* infection induced a non-significant decrease in BLOC1S1 expression in all cell types which was exacerbated by a productive HIV-1 pre-infection in glial cells (Supplementary Figure 4E-H).

Similar to spliced XBP1, a significant increase in CHOP transcript following *Mtb* infection occurred in astrocytes (1.95-fold), endothelial cells (2.18-fold), and pericytes: (4.07-fold, Fig. 3I-L). Co-infection with HIV-1 slightly exacerbated the increase in endothelial cells and pericytes. The only significant change in transcript of BiP was a small *Mtb-*induced decrease in the presence of HIV-1_BAl_ in astrocytes (Supplementary Figure 4M-P). *Mtb* infection also induced a small decrease (1.14-fold) of XBP1 in astrocytes, and small increases in endothelial cells, pericytes and microglia (1.57-fold, 1.86-fold, and 1.17-fold, respectively). Co-infection with productive HIV-1 tended to augment the *Mtb*-induced decrease in XBP1 expression in astrocytes.

### Effect of co-infection on astrogliosis

We next investigated the effect of infection on astrogliosis by measurement of the expression of complement 3 (C3, a neurotoxic reactive astrocyte marker) and S100A10 (a neuroprotective marker). *Mtb* infection induced a 40.36-fold increase in C3 expression with HIV-1_Bal_ infection causing a further slight increase (2.24-fold median increase compared to *Mtb* infection, Fig.5A). By contrast, S100A10 expression was not significantly changed by either infection (Fig. 5B).

### Effect of *Mtb* infection on glutamate concentration

The homeostasis of the neurotransmitter glutamate in the brain extracellular space is tightly regulated^11–13^. Because *Mtb* and HIV-1 infections caused CNS cell stress, we investigated the effect of both infections on extracellular glutamate both in single cell monolayer (2 dpi) and in our BBB model (2 and 6 dpi) (Fig. 5C-H). Neither route of HIV-1 infection alone affected extracellular glutamate in either the BBB model, or in isolated cells. By contrast, *Mtb* infection continuously increased extracellular glutamate 1.4-fold at 2, and 3.8-fold at 6 dpi compared to uninfected BBB control cultures. No additional effect of HIV-1 infection was observed (Fig. 5C,D). *Mtb* also significantly increased extracellular glutamate in cultures of astrocytes and microglia especially (1.8- and 1.7-fold, Fig. 5E,H) and to a lesser extent pericytes (1.2-fold, Fig. 5G). Endothelial cells did not appear to contribute to extracellular glutamate homeostasis (Fig. 5F). Again, no significant additional effects were observed in the presence of HIV-1 infection.

### Effect of *Mtb* and HIV-1 infections on soluble mediator secretion

The inflammatory response of the host contributes to TBM pathology^14^. Thus, we determined the effect of both *Mtb* and HIV-1 infections on the release of matrix metalloproteinases (MMP), vascular endothelial growth factors (VEGF), interleukins, and chemokines from CNS cells and in our BBB model.

VEGF-D, IL-12p70, IL-10, IL-13, IL-17A, and eotaxin-2 were not detected (median 0 pg/ml), whilst infection-induced MMP-9, eotaxin-1 and −3, IL-2, and IFN-γ by CNS cells above background was negligible or even decreased. Low but detectable increases in IL-1α, IL-4, MIP-1α, MIP-1β, TNF, fractalkine and RANTES occurred upon *Mtb* infection and *Mtb*/HIV-1 co-infection, depending of the cell type (Table S1).

Principal component analysis (PCA) of the data revealed that cell types clustered distinctly, indicating that the effector cell type was an important determinant of the response (Fig. 6A). Figure 6B shows only analytes with consistently detectable median values and indicates a dominant effect of *Mtb* over HIV-1 infection. Constitutive MMP-2 and MCP-1 secretions tended to decrease in the BBB and all cell types, although, interestingly, *Mtb*/VSV-G pseudotyped HIV-1 co-infection induced an increase of MMP-2 concentration in microglia (3.1- and 6.2-fold increase compared to uninfected and *Mtb* mono-infected conditions, respectively). Constitutive MMP-3 secretion decreased 1.3-fold during *Mtb* infection in pericytes and increased 2.5-fold in microglia. These changes were modestly accentuated by HIV-1 co-infection.

**Figure 6.**
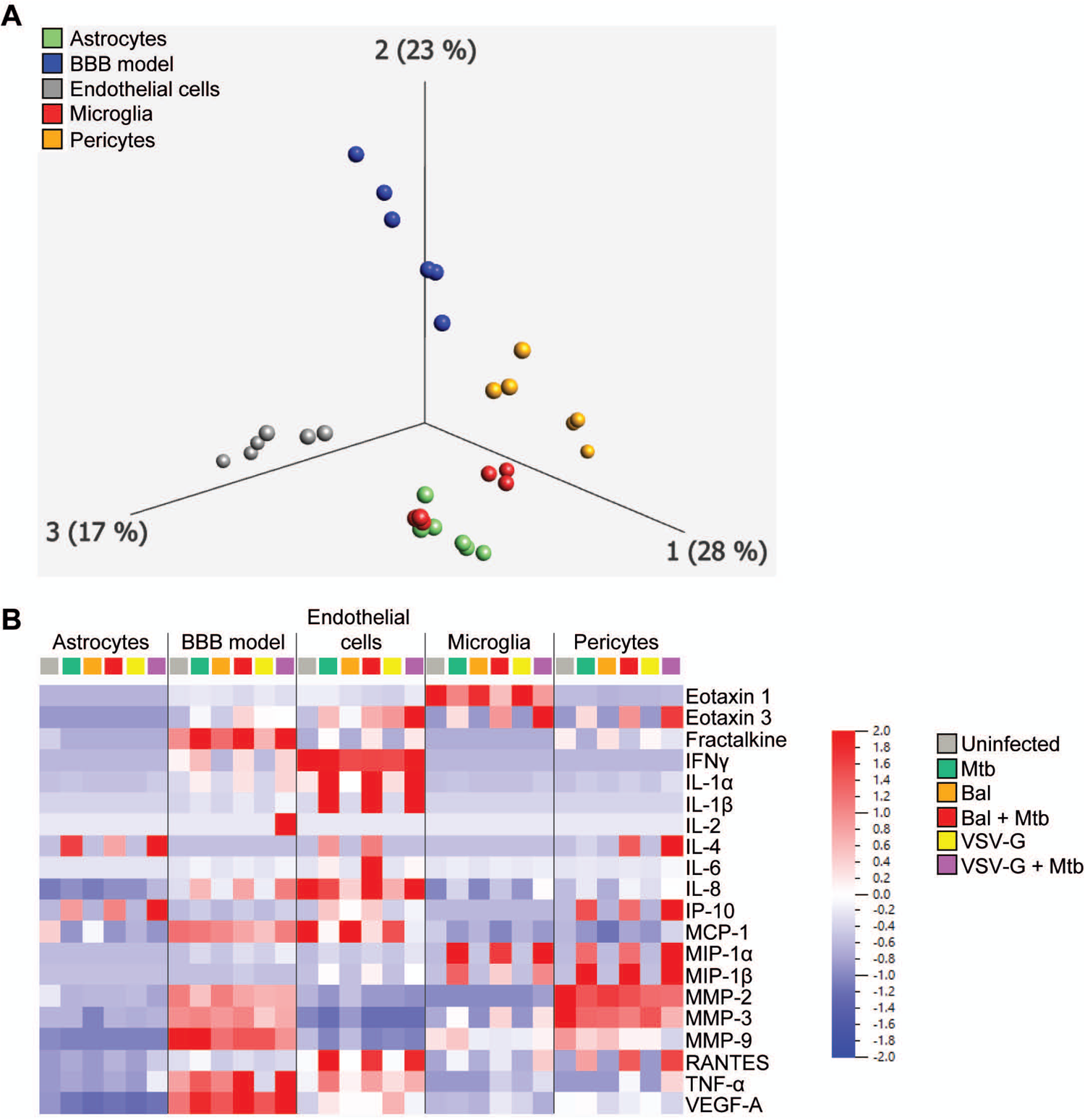
CNS cell and BBB soluble factors released in response to *Mtb* +/− HIV-1 infection (s) (A) Principal component analysis clustering astrocytes (green), endothelial cells (grey), microglia (red), pericytes (orange) and BBB model responses (blue). (B) Normalised heatmap of analytes with detectable median values.

Secretion of IL-1β by endothelial cells was increased by *Mtb* infection (102.3-fold increase compared to uninfected), reflected in the BBB model with IL-1β concentrations increasing from 0 pg/ml to 5.21 pg/ml upon *Mtb* infection. HIV-1 infection did not modulate IL-1β secretion by endothelial cells, but co-infection increased its secretion in the BBB model (1.4- and 1.8-fold increase compared to *Mtb* mono-infection for *Mtb*/Bal and *Mtb*/VSV-G co-infections, respectively). *Mtb* infection increased IL-6 and IL-8 secretion in most cell types (except IL-8 from endothelial cells) and in the BBB model. These increases were modestly and variably accentuated by both forms of HIV-1 co-infection. Although *Mtb* decreased VEGF-A secretion 1.5-fold by astrocytes, it increased secretion from endothelial cells and in the BBB model with *Mtb*/HIV-1 co-infection slightly accentuating VEGF-A secretion in the BBB model. *Mtb* variably increased secretion of IP-10 from astrocytes, endothelial cells, pericytes, and the BBB model and productive HIV-1 co-infection accentuated release in astrocytes (2.4-fold), pericytes (1.7-fold) and the BBB model (1.3-fold).

## Discussion

Dissemination of *Mycobacterium tuberculosis* to the brain results in the most severe form of extrapulmonary TB, tuberculous meningitis, which constitutes a medical emergency associated with elevated mortality and disability. In this study, we investigated effects of *Mtb* on BBB and CNS cells to gain better understanding of *Mtb* spread to the brain and the influence of HIV-1 on that process. A summary of our findings is shown in Supplementary Table 2.

While it is recognised that *Mtb* disseminates into the CNS^7^, little is known about the susceptibility of CNS cells to *Mtb* infection. Microglia are susceptible to *Mtb* and have been proposed as the main target cells of *Mtb* in the CNS^15^. Jain and colleagues described that brain microvascular endothelial cells can be invaded by both laboratory H37Rv and clinical CDC1551 *Mtb* strains^9^. By contrast Rock and colleagues showed astrocytes were moderately susceptible to infection^15^. We observed that *Mtb* enters and replicates in CNS cells, and HIV-1 increases entry to astrocytes and pericytes, and growth in pericytes and endothelial cells. Length of infection may influence differences between our results (around 40% of astrocytes were infected after 2 days at MOI 5) and those of Rock (15% of astrocytes were infected after 24 hrs at MOI 10). Interestingly while *Mtb* preferentially entered HIV-1 uninfected cells, it replicated better in HIV-1 productively infected cells.

The impact of *Mtb* on the BBB and whether it disseminates to the brain as cell free bacilli or via a Trojan horse mechanism with cells is still debated^16^. Using a 3D BBB model, we showed that *Mtb* steadily decreases BBB integrity over time, congruent with its cytopathic effects. While *Mtb*-induced BBB permeability was only slightly increased by HIV-1, the concentration of bacilli translocated through the model after two days of infection was significantly increased by bystander HIV-1 co-infection. This suggests HIV-1 facilitates *Mtb* transmigration through the BBB and into the brain.

Neither *Mtb* infection nor *Mtb*/HIV-1 co-infection significantly affected the expression of tight junction proteins although there was a slight decrease in the expression of VE-cadherin, an adherens junction essential for endothelium integrity, endothelial cell survival and angiogenesis^17^ (Figure S2). Overall, these results suggest that (i) *Mtb* can disseminate from the blood to the brain directly rather than within cells, (ii) HIV-1 facilitates *Mtb* transmigration through the BBB during the early phase of infection, (iii) *Mtb* causes a loss of BBB integrity although does not downregulate gene expression for tight junction proteins expressed by endothelial cells, (iv) *Mtb* transmigration therefore could be paracellular (through a leaky BBB) but (v) a transcellular transmigration (through cells of the BBB) appears operative as *Mtb* rapidly entered and grew in CNS cells.

Mitochondrial metabolism, ROS production and ER stress are closely linked and involved in a variety of physiological and pathological conditions. Mitochondrial function is susceptible to pathological insults leading to ER stress. Depending on the degree of cellular stress, the signalling from the ER to mitochondria can either lead to pro-survival or pro-apoptotic adaptations of mitochondrial function. Early during ER stress, mitochondrial metabolism is stimulated due to increased Ca^2+^ transfer between ER and mitochondria. Later ER stress phases are characterised by decreased mitochondrial respiration, negatively affecting cellular metabolism^18, 19^. In parallel, ROS can be generated by both stressed mitochondria and endoplasmic reticulum and their production has been suggested as integral UPR component^20^. We showed that *Mtb* infection has no (astrocytes and pericytes) or little (endothelial cells and microglia) impact on mitochondrial metabolism. ROS production is involved in the pathogenesis of both TB and TBM^21^. We showed that *Mtb* triggered ROS generation by CNS cells could be a host defence mechanism as in macrophages^22^. HIV-1 co-infection had, however, no effect on ROS production which differs from a previous report^23^. As we measured H_2_O_2_ levels induced by full-length live virus, the discrepancy may be explained by the type of cells^24^ or the type of infection (full virus/viral components/transfection)^25–27^.

Both HIV-1 and *Mtb* infections trigger the UPR^28, 29^. Tripathi and colleagues showed elevated levels of CHOP, ATF4 and BiP in the blood of TBM patients *versus* healthy individuals and in severe TBM *versus* mild TBM patients, suggesting that ER stress relates to the severity of infection^30^. Hitherto there were no data on *Mtb* induced ER stress in CNS cells. Our study tends to confirm previous results obtained from blood cells. We showed that *Mtb* activates the IRE1α and PERK branches of the UPR in CNS cells. Both pathways participate in the induction of autophagy during ER stress, which precedes induction of apoptosis induction which is seen as beneficial for cell survival^31^. However, if the acute ER stress (adaptive UPR) fails to restore protein-folding homeostasis, the UPR signalling switches into chronic ER stress (terminal UPR) that ultimately promotes apoptosis. This fate-switch is regulated by IRE1α (transient sXBP1 expression induces autophagy while sustained sXBP1 levels lead to apoptosis) and PERK (CHOP overexpression results in apoptosis and/or cell arrest) pathways^32^. Sustained expression of CHOP and high levels of sXBP1 in *Mtb*-infected CNS cells suggest that the cells reach a terminal UPR state upon *Mtb* infection.

Astrogliosis represents a spectrum of changes in astrocytes occurring in response to neuronal insults and is promoted by various cytokines including IL-6, TNF-α and IL-8^33^. Upon brain insult, astrocytes can either become neuroprotective (A2 reactive astrocytes) and exert beneficial functions (restriction of CNS inflammation, synapses repair, neuronal protection, BBB repair, and wound closure) or neurotoxic (A1 astrocytes) that exacerbate inflammation and interfere with synapse sprouting or axonal growth^34^. Reactive astrocytes express specific markers (S100A10 for A2 astrocytes, complement 3 for A1 astrocytes) allowing their identification^35, 36^. We demonstrated that not only both *Mtb* and HIV-1 productive infections induce A1 astrocytes but also that co-infection with HIV-1_Bal_ worsens the *Mtb*-induced effect, suggesting a bystander effect of HIV-1 on astrogliosis.

Glutamate is the main excitatory neurotransmitter in the mammalian CNS where its concentration is tightly regulated to prevent both excitotoxicity and defective neurotransmission^13^.Extracellular glutamate levels were increased in *Mtb*-infected astrocytes, pericytes, microglia and our BBB model but were not worsened by HIV-1 co-infection. Elevated levels of glutamate and transcripts associated with glutamate release have been measured in the CSF of paediatric TBM patients^37, 38^ which accords with our results. In a healthy CNS, astrocytes are the main cells maintaining glutamate homeostasis, taking up glutamate via the glutamate transporter 1 (GLT-1). In a pathological CNS, astrocytic glutamate uptake is impaired by metabolic stress, ROS and HIV-1 infection^39–41^. Activated microglia can also release glutamate, alter gliotransmission and interfere with glutamate uptake by astrocytes^12^. Endothelial cells, capable of uptaking and metabolising glutamate, can also play a role in glutamate homeostasis^11^. Our results suggest infection of CNS cells and their release of ROS upon *Mtb* infection could explain this increase in glutamate levels.

A meta-analysis study reported elevated CSF levels of IL-1β, IL-2, IL-4, IL-6, IL-8, IL-10, IFN-γ and TNF-α in TBM patients compared to healthy individuals^42^. The same analysis showed that IL-1β, TNFα and IL-2 were further increased in HIV-positive persons with TBM *versus* HIV-negative while their levels of IFN-γ, IL-10 and IL-12p70 were decreased. Visser and colleagues also showed increased CSF concentrations of IL-1β, IL-6, IL-12, IFN-γ, TNF, IL-10, IL-13, IP-10, IL-8, MIP-1α, MIP-1β, RANTES and VEGF in TBM patients^43^. Both findings are concordant with ours. We showed that *Mtb* induces a neuroinflammatory environment supporting a loss of BBB integrity. It is of note that in our study, apart from MMP-2, MMP-3, IL-6, IL-8, VEGF-A and MCP-1, soluble factors levels detected were low, consistent with the absence of peripheral immune cells in the model.

In conclusion *Mtb* infects and multiplies in CNS cells with HIV-1 increasing entry to astrocytes and pericytes, and growth in HIV-1 infected pericytes and endothelial cells. The permeability of the BBB increases resulting in translocation of bacilli across it. These data suggest that *Mtb* can translocate the BBB directly and initiate an immune response that leads to further cellular recruitment that initiates an inflammatory meningitis.

## Online Methods

### Ethical approval

Human foetal brain tissues from 15 to 20 weeks’ foetuses were obtained from the MRC-Wellcome Trust Human Developmental Biology Resource (HDBR), UCL, with ethical approval (University College London, UCL, site REC reference: 18/LO/0822 - IRAS project ID: 244325 and Newcastle site REC reference: 18/NE/0290 - IRAS project ID: 250012).

### Commercial human cells and culture conditions

Human cerebral microvascular endothelial cells (hCMEC/D3, Merck Life Science) were grown on collagen I-coated (150 µg/mL, Sigma-Aldrich) flasks/wells using the EndoGRO^TM^-MV complete Media Kit supplemented with 1ng/mL FGF-2 (“hCMEC/d3 medium”, both from Merck Life Science). Human microglial cells (HMC3, ATCC) were grown in EMEM (ATCC) supplemented with 10 % heat-inactivated foetal bovine serum (Corning). Cell lines were used after 3-12 passages throughout. Human Brain Vascular Pericytes (HBVP, ScienCell) were grown on Poly-L-Lysine (2 µg/cm^2^, Sigma-Aldrich) coated flasks/wells in complete Pericyte Medium (ScienCell). All cells were grown at 37°C in the presence of 5% CO_2_.

### Primary human astrocyte isolation

Human foetal astrocytes were isolated from foetal brain samples as previously described^44^. Briefly, blood vessels and meninges were removed from the foetal brain tissue (15 to 20 gestational weeks). Thereafter, the tissue was minced, treated with 0.2 mg/ml DNase I (Sigma-Aldrich) and 0.25% trypsin (ThermoFisher Scientific) for 30 min before being passed through a 70-µm cell strainer (Corning). The flow-through was plated in petri dishes for adherent cells (Sarstedt) at a final concentration of 6-8 × 10^7^ cells/petri in MEM supplemented with 10% FBS, 100 U/ml penicillin, 100 µg/ml streptomycin, 0.3mg/ml L-glutamine, 1 mM sodium pyruvate, 1X MEM nonessential amino acids, 0.5 µg/ml amphotericin B, and 2.5ml of glucose solution (all from ThermoFisher Scientific). Astrocytes were grown at 37°C (5% CO_2_) and left undisturbed for two weeks, thereafter they were passaged once every week. To ensure cell purity, all experiments were conducted using cells after 3 to 6 passages.

### Bacterial strain, growth condition and preparation for cell infection

We used an RFP fluorescent *Mtb* was obtained from Maximiliano G. Gutierrez^45^. The fluorescent *Mtb* H37Rv is engineered to constitutively express the Red fluorescent protein (RFP) encoded by pML2570 plasmid and integrated into the bacterial genome (integrase: Giles, resistance: Hygromycin). H37Rv-RFP were grown in Middlebrook 7H9 broth supplemented with 0.5% glycerol (Fisher Chemical), 0.05% Tween-80 (MP Biomedicals), 10% ADC (Sigma-Aldrich) and 50 µg/mL Hygromycin (Invitrogen, Carlsbad, CA). Bacterial cultures were incubated at 37 °C with rotation in 50 mL conical tubes. To infect cells, *Mtb* was grown to an OD_600_ of 0.5 to 1.0 and washed 3 times (twice in PBS and once in cell culture medium, 5 minutes at 3500 rpm). The bacteria pellet was resuspended in 10 mL cell culture medium and shaken with 2.5–3.5 mm sterile glass beads for 1 min to produce a single-cell suspension. The OD_600_ was determined, and bacteria diluted for the required multiplicity of infection (MOI), assuming OD_600_ 1 = 1×10^8^ bacteria/ml.

### Virus production

Virions were produced by calcium-phosphate transient transfection of 293T cells as previously described^46^. HIV-1 Bal virus was produced by transfecting 293T with the NL4-3-Bal-IRES-HSA plasmid^47^. VSV-G-pseudotyped HIV-1 virus was produced by co-transfecting 293T with pHCMV-G and NL4-3-IRES-HSA *env-* plasmids. Expression vectors were kindly provided by Dr Michel J. Tremblay (Centre de Recherche en Infectiologie de l’Université Laval, Canada). Infectivity of the virus stocks was assessed using the genetically modified HeLa-derived indicator cell line TZM-bl and the Spearman-Karber method^46^.

### *In vitro* human BBB model system

We used a co-cultivation model including hCMEC/D3 cells and a mix of pericytes and astrocytes seeded on each side of a porous insert allowing cell-cell contact as previously described ^48^. Microglia were seeded on the bottom of the well to mimic their localization in the CNS. Briefly, cell culture inserts for 24-well plates with a 3.0-μm pore size, translucent PET membrane (Corning) were coated with 150 μg/mL collagen-I (Sigma-Aldrich) on the upper side and with Poly-L-Lysine (2 µg/cm^2^, Sigma-Aldrich) on the basal side. A mix of HBVP (10^4^/insert) and astrocytes (5×10^4^/insert) were then seeded on the basal side of the membrane and incubated for 4 h before hCMEC/D3 cells (2.5×10^4^/insert) were seeded on the upper side of the membrane. Cells were allowed to grow in 150 μL (upper chamber) and 750 μL (collector) of BBB medium 1, composed of 50% hCMEC/d3 medium and 50% HBVP medium, for 7 days to reach confluence (medium in both upper chamber and collector was refreshed with 150 μL and 750 μL of BBB medium 1, respectively, at day 4). HCM3 (5×10^3^/well) were then seeded at the bottom of the well containing the inserts and medium changed (upper chamber and collector) to a mix of 50% hCMEC/d3 medium, 25% HBVP medium and 25% HMC3 medium (BBB medium). BBB integrity was assessed by measuring permeability to dextran-rhodamine B. Briefly, the culture medium in the upper chamber was replaced with BBB medium supplemented with 0.5 mg/mL 70-kDa dextran-rhodamine B (Life Technologies). After 5h, the fluorescence intensity in the collector was measured using a Synergy 2 multi-mode microplate reader (Biotek). Samples displaying dextran-rhodamine B permeability > 20% in the empty control insert were discarded.

### CNS cell infection with HIV-1 and/or *Mtb*

CNS cells seeded either on inserts (BBB model), E-plates (xCELLigence assay), 96-well plates (MTS, ROS and glutamate assays – 2.5×10^4^ cells/well), 24-well plates (*Mtb* entry and growth assays) or 12-well plates (quantification of mRNA - 2×10^5^ cells/well) were either infected with HIV-1 (Bal or VSV-G, MOI 0.1) or left uninfected for 24 hrs. before being extensively washed to remove unbound viral particles. Cells were then either infected with *Mtb* (MOI 5) or left uninfected for 3 hrs before extensive washes. Cells were then left untouched for 2 or 6 days.

### *Mtb* entry and growth in CNS cells

Cells seeded in 24-well plates (1×10^5^ cells/well) were infected with HIV-1 +/− *Mtb* as described above. For both *Mtb* entry and growth experiments, cells were extensively washed after the 3 hrs of incubation with *Mtb* and then either detached directly (*Mtb* entry) or after 48 hrs (*Mtb* growth) using TrypLE^TM^ Express 1X (Gibco). Cells were then stained with eFluor^TM^450 fixable viability dye (Invitrogen) for 30 min at room temperature (RT), blocked with Pharmingen^TM^ stain buffer BSA (BD Bioscences) for 30 min at RT and stained with an FITC-conjugated anti-CD24 monoclonal antibody (Life Technologies) for 20 min at RT. Stained cells were then fixed with 4% paraformaldehyde overnight and permeabilized before acquisition on a LSRFortessa^TM^ cell analyzer (BD Bioscences). Data were analysed with FlowJo software v10.9.0.

### BBB permeability assay

The effect of *Mtb* and/or HIV-1 infections on BBB integrity was assessed by measuring the permeability of the BBB to dextran-rhodamine B. After 2 days or 6 days following *Mtb* infection the culture medium in the insert was replaced with BBB medium supplemented with 0.5 mg/mL 70-kDa dextran-rhodamine B. The fluorescence intensity in the collector was measured using a Synergy 2 multi-mode microplate reader (Biotek) after 5 hrs.

### Quantification of *Mtb* passage through the BBB model

The concentration of viable bacteria passing through the BBB model was determined by colony forming unit assay (CFU). BBB model collectors after 48 hrs. of *Mtb* infection were centrifuged at 3500 rpm for 5 minutes to concentrate the bacteria 15 fold. Concentrated collectors were plated in triplicate onto complete 7H11 agar trisector plates supplemented with 50 µg/mL Hygromycin (25 µL/sector). Agar plates were incubated then for 2–3 week at 37°C, the number of colonies per sector counted and CFU calculated.

### xCELLigence assay

The xCELLigence Real Time Cell Analysis Dual Purpose system (RTCA DP, Agilent) allows the continuous and non-invasive recording of electrode impedance integrated into the bottom of E-plates 16. The number, size and shape of cells affects the impedance which is reported using the unitless parameter cell index (CI). The CI is directly proportional to the relative change in electrical impedance normalized by the background value. Recording of CI values and analysis were performed using the Real Time Cell Analysis Software Pro version 2.3 (Agilent). In brief, hCMEC/D3 cells, astrocytes, HMC3 and HBVP were evenly distributed in 16-well E plates pre-coated with Collagen-I (hCMEC/d3 cells) or Poly-L-Lysine (astrocytes and HBVP) in three to four replicates for each condition. To prevent edge effects, we allowed the cells to attach to the 16-well E-plates at room temperature for 30 min before being inserted in the RTCA DP system, inside an incubator (37°C, 5% CO_2_) for continuous impedance recording. The hCMEC/D3 cells reached their confluent phase and formed a tight barrier on day 4-5. Astrocytes, HMC3 and HBVP were confluent on the day following their seeding. Cells were then infected by HIV-1 and/or *Mtb* as described above. The uninfected condition was used as baseline and negative control in our analysis.

### Tight and adherens junction expression

hCMEC/d3 cells were seeded on 12-well plates pre-coated with Collagen-I for 6 days before infection as previously described, and mRNA quantification.

### Quantification of mRNA

Total RNA was extracted after 48 hrs. of *Mtb* infection using TRIzol^TM^ reagent (Invitrogen) and the Zymo Direct-zol 96-RNA kit (Zymo Research) according to the manufacturer’s instructions, followed by reverse transcription to cDNA with M-MLV RT Polymerase (Promega). Gene expression was assessed by SYBR-Green Quantitative RT-PCR using QuantStudio^TM^ 7 Flex Real-Time PCR System (Thermo Fisher Scientific) following the manufacturer’s instructions. Primers were used at 0.3 μM (Supplementary Table 3). Amplification of target genes was normalized to the geometric mean of 18S ribosomal cDNA. A standard curve was drawn for each gene of interest using serial dilutions of pooled cDNA from all samples.

### MTS assay

After 48 hrs *Mtb* infection, the metabolic activity of CNS cells was assessed using CellTiter 96™ AQueous Nonradioactive Cell Proliferation Assay (conversion of the tetrazolium salt MTS [3-(4,5-dimethylthiazol-2-yl)-5-(3-carboxymethoxyphenyl)-2-(4-sulfophenyl)-2H-tetrazolium)] to a purple formazan in the presence of phenazine ethosulfate) following the manufacturer’s instructions (Promega). Absorbance at 490 nm was measured using a Synergy 2 multi-mode microplate reader (Biotek instruments).

### ROS assay

The level of hydrogen peroxide (H_2_O_2_) after 48 hrs. infection with *Mtb* was measured using the ROS-Glo^TM^ H_2_0_2_ assay following the manufacturer’s instructions (Promega). Menadione at 50 uM was used as positive control. Luminescence was measured using a Synergy 2 multi-mode microplate reader (Biotek instruments).

### Glutamate assay

Glutamate levels after 48 hrs. of *Mtb* infection were monitored using the Glutamate assay kit (Abcam) on cell experiment supernatants and BBB collectors following the manufacturer’s instructions. Optical density at 450 nm was measured using a Synergy 2 multi-mode microplate reader (Biotek instruments).

### Multiplex immune assay for cytokines and chemokines

Soluble factors secreted by CNS cells in response to *Mtb* +/− HIV-1 infection(s) were detected in the cell experiment supernatants and BBB collectors by ProcartaPlex^TM^ immunoassay (Life Technologies) using a Bio-Plex 200 system (Bio-Rad, Hercules, CA) according to manufacturer’s instructions. Visualisation of the results was performed using Qlucore Omics Explorer v.3.3 (Qlucore).

### Statistical analysis

Statistical analysis was performed using GraphPad Prism version 10.0.3. Statistical details of each experiment (statistical tests used, exact value of n, dispersion, and precision measures) can be found in the figure legends. Means ± SEM or median ± IQR were used according to the normality of data throughout. Statistical analyses were performed on raw data for all experiments. A threshold p value of ≤ 0.05 (*) was considered statistically significant. When non statistically significant, no stars have been added in figures for the sake of clarity.

## Acknowledgements

The authors thank Michel J. Tremblay and his team at the CHU de Québec-Université Laval research centre for providing NL4-3-Bal-IRES-HAS, pHCMV-G and NL4-3-IRES-HSA *env-* plasmids. We also thank the Flow Cytometry platform of the Francis Crick Institute. The human foetal material was provided by the Joint MRC/Wellcome Human Developmental Biology Resource (www.hdbr.org) (project#200511). The authors received support from the Francis Crick Institute which is funded by Wellcome (CC2112), Cancer Research UK (CC2112) and UK Research and Innovation, Medical Research Council (CC2112). Additional funding was provided by NIH (R01AI145436) and Meningitis Now. For the purposes of open access, the authors have applied a CC-BY public copyright to any author-accepted manuscript arising from this submission.

## Author’s contributions

AP: conceptualization, formal analysis, investigation (lead), methodology, project administration, validation, and writing of original draft. KAW: investigation (supporting), writing, review and editing (supporting). RJW: funding acquisition, supervision, writing, review and editing (lead).

## Data availability statement

Raw data from flow cytometry (figures 1 and S1), CFU counts and dextran rhodamine B fluorescence readings (figure 2), real-time cell analysis with xCELLigence (figure 3), MTS and ROS assays (figures 4 and S3), astrogliosis qPCR and glutamate fluorescence readings (figure 5), Luminex (figure 6 and table S1), tight junction and ER stress qPCR are publicly available in the Crick figshare repository (10.25418/crick.26869453)

## Notes

### Competing Interest Statement

The authors have declared no competing interest.

