## Supplementary information for "M. tuberculosis invades and disrupts the blood brain barrier directly to initiate meningitis"

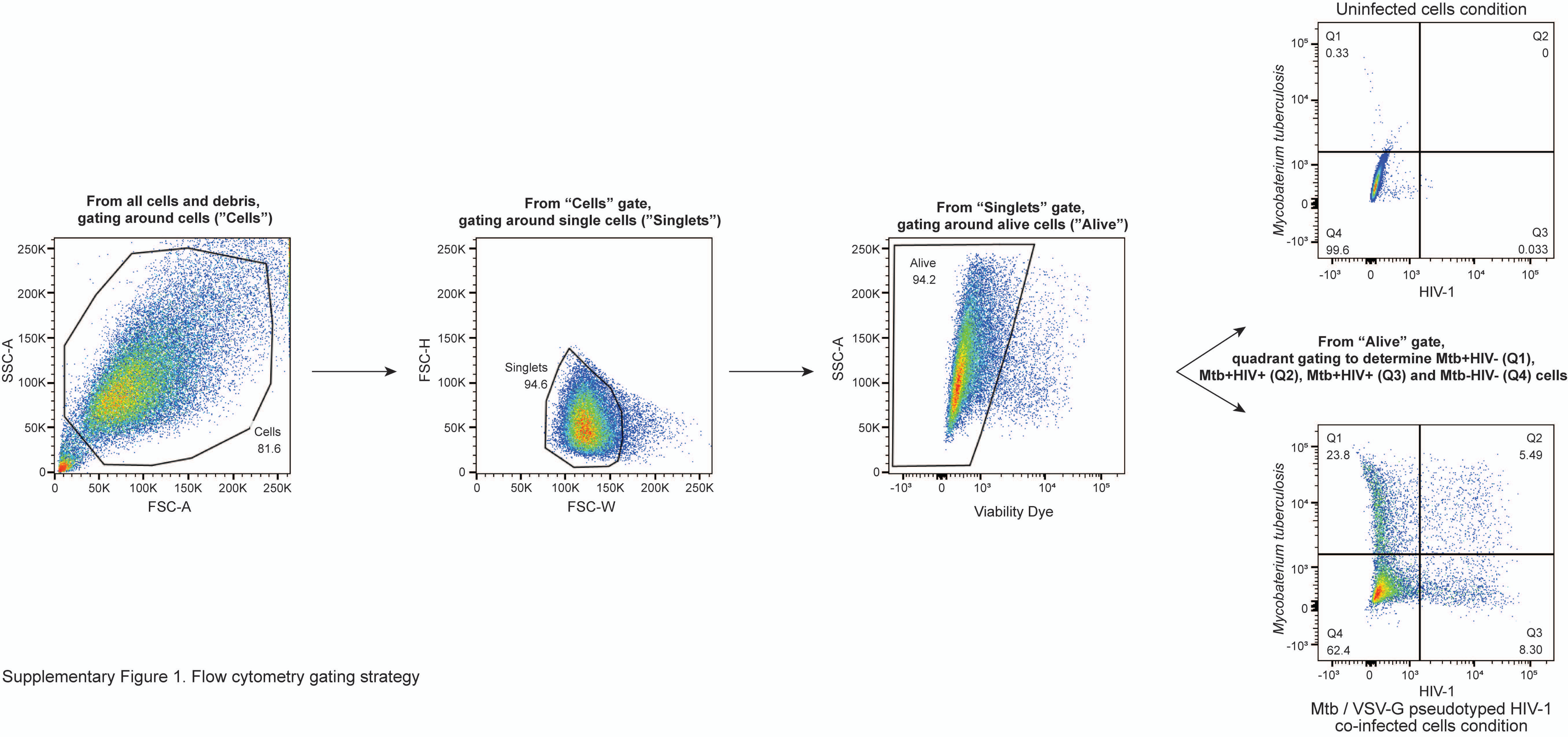


Supplementary Figure 1. Flow cytometry gating strategy

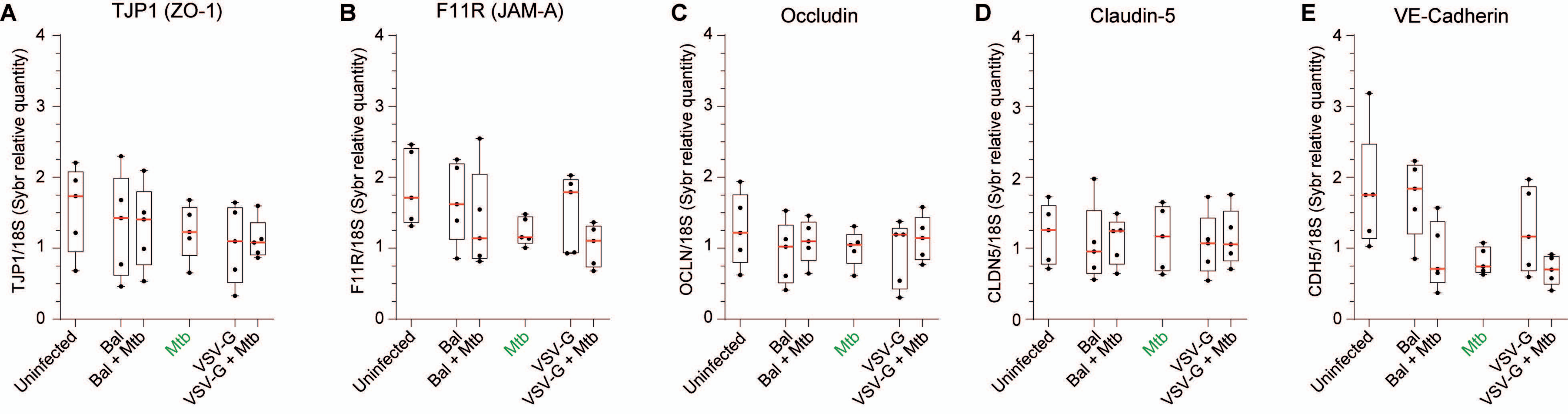


Supplementary Figure 2. Effect of Mtb and HIV-1 infection on endothelial cell tight cell, and adherens junction gene expression.

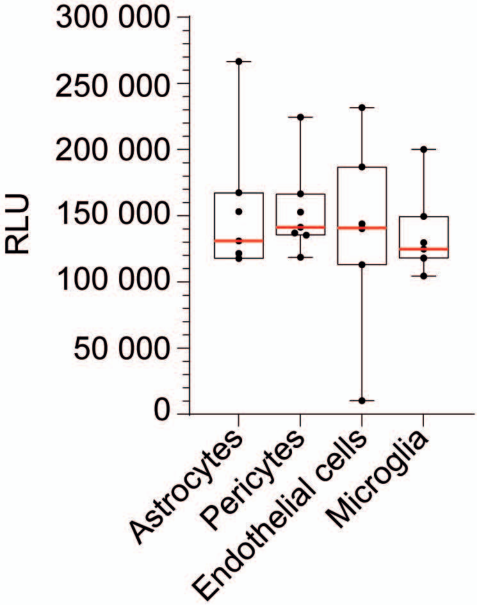


Supplementary Figure 3. ROS assay positive control.

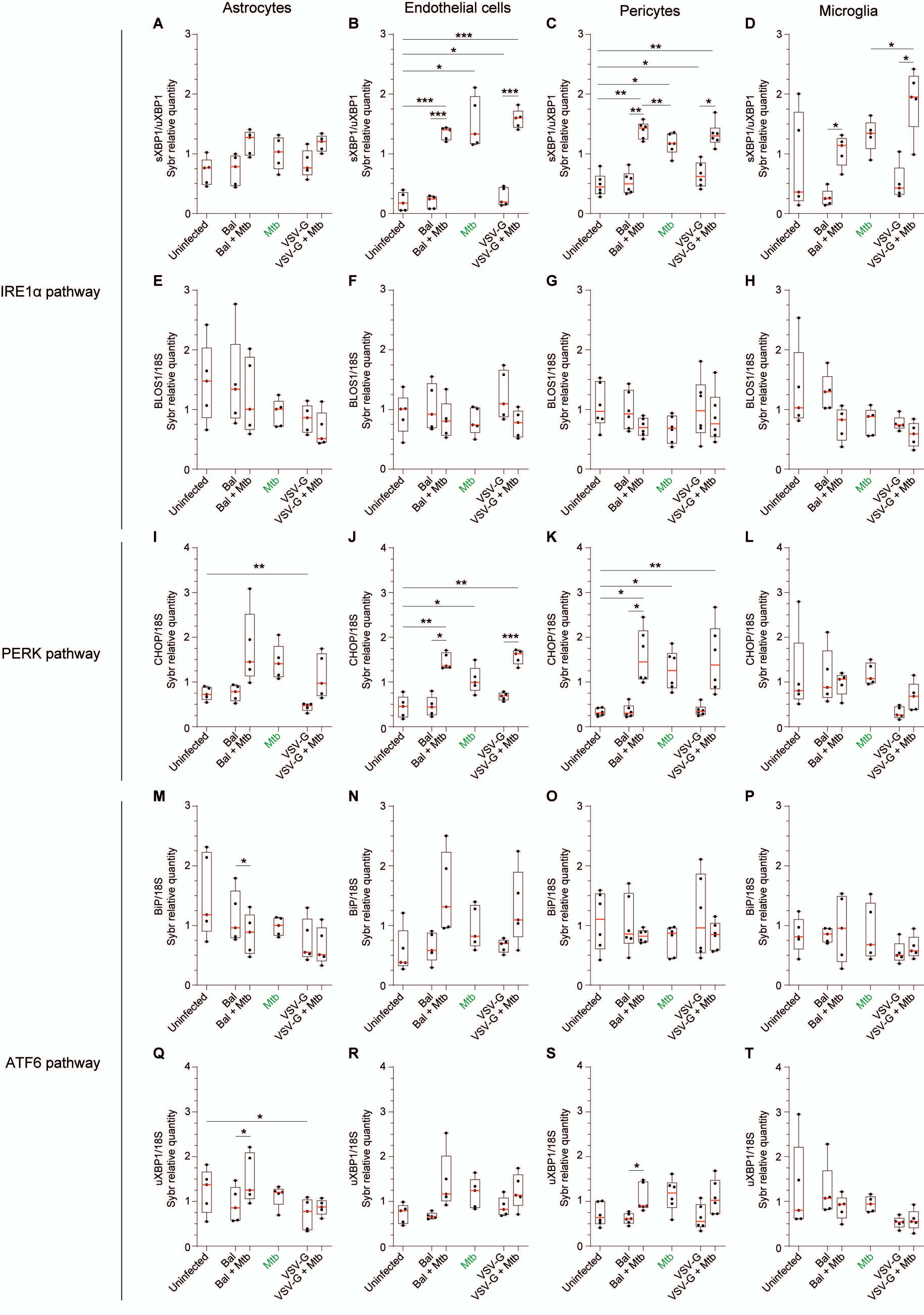


Supplementary Figure 4. Effect of HIV-1 and/or Mtb infection(s) on ER stress.

| | | MMPs | | | Eotaxins | | | Interleukins | | | | | | | | | | MIPs | | VEGFs | | Fractalkine | IFNγ | TNFα | IP-10 | MCP-1 | RANTES |
|---|---|---|---|---|---|---|---|---|---|---|---|---|---|---|---|---|---|---|---|---|---|---|---|---|---|---|---|
| | | 2 | 3 | 9 | 1 | 2 | 3 | 1α | 1β | 2 | 4 | 6 | 8 | 10 | 12p70 | 13 | 17A | 1α | 1β | A | D | | | | | | |
| Astrocytes | Uninfected | 841.51 (433.07-1222.53) | 47.18 (23.72-63.22) | 0.27 (0-1.56) | 0.03 (0-0.28) | 0 | 0 | 0 | 0 | 0 | 0 | 51.31 (33.24-147.64) | 13.18 (5.74-63.05) | 0 | 0 | 0 | 0 | 0 | 0 | 822.64 (445.15-1258.98) | 0 | 2.09 (0-7.54) | 0 | 0 (0-0.38) | 0.45 (0-1.11) | 1023.53 (879.55-1519.51) | 0 (0-1.32) |
| | Mtb | 386.48 (190.17-764.48) | 46.47 (9.03-65.92) | 0 | 0 (0-0.16) | 0 | 0 | 0.14 (0-0.25) | 0 | 0 | 11.12 (0.35-15.07) | 333.71 (252.69-542.69) | 423.67 (106.77-629.9) | 0 | 0 | 0 | 0 | 0 | 0 | 545.84 (393.66-939.15) | 0 | 0 (0-4.18) | 0 | 0.03 (0-0.2) | 85.81 46.27-180.93) | 424.58 (368.53-503.66) | 2.21 (0-5.4) |
| | HIV-1 Bal | 412.33 (251.54-1160.12) | 15.73 (0-42.96) | 0 | 0 | 0 | 0 | 0 | 0 | 0 | 0 | 64.26 (0-128.74) | 2.09 (0-22.21) | 0 | 0 | 0 | 0 | 0 | 0 | 416.01 (337.46-1070.15) | 0 | 0 (0-6.99) | 0 | 0 (0-0.32) | 0 (0-0.32) | 797.33 (629.96-1033.57) | 0 (0-1.01) |
| | Bal + Mtb | 411.81 (251.54-1160.12) | 46.47 (0-64.7) | 0 | 0 | 0 | 0 | 0.01 (0-0.16) | 0 | 0 | 6.69 (5.21-12.84) | 380.36 (273.23-1795.42) | 439.78 (80.92-801.47) | 0 | 0 | 0 | 0 | 0 (0-2.18) | 0 | 502.91 (288.78-917.96) | 0 | 0 (0-3.38) | 0 | 0.06 (0-0.5) | 97.83 (28.66-174.37) | 411.16 (381.29-456.56) | 3.12 (0.28-9.66) |
| | HIV-1 VSV-G | 462.56 (182.46-764.48) | 44.48 (0-73.12) | 0 (0-0.1) | 0 | 0 | 0 | 0 | 0 | 0 | 0 | 167.89 (68.90-345.1) | 361.91 (66.2-819.31) | 0 | 0 | 0 | 0 | 0 | 0 | 428.6 (216.89-717.31) | 0 | 0 (0-2.49) | 0 | 0 (0-0.46) | 0.19 (0-4.12) | 530.59 (549.91-696.46) | 0.42 (0-1.32) |
| | VSV-G + Mtb | 244.39 (190.9-653) | 48.46 (22.5-88.21) | 0 (0-0.1) | 0 (0-0.04) | 0 | 0 (0-0.38) | 0.33 (0-0.6) | 0 | 0 | 17.14 (9.97-32.53) | 367.32 (276.99-1853.56) | 1187.24 (240.37-1531.24) | 0 | 0 | 0 | 0 | 0 (0-1.91) | 0 | 524.84 (391.66-826.38) | 0 | 0 | 0 | 0.38 (0.1-1.12) | 205.88 (90.08-222.72) | 420.23 (350.80-483.67) | 6.87 (3.35-16.98) |
| Endothelial cells | Uninfected | 381.45 (114.43-1005.87) | 17.58 (5.77-74.89) | 1.52 (0.1-2.69) | 1.57 (0.54-1.7) | 0 | 0.09 (0-0.21) | 2.48 (1.3-5.36) | 1.64 (0.95-3.92) | 0 (0-5.35) | 0 | 275.06 (79.27-353.88) | 7506.0 (2680.27-10043.42) | 0 | 0 (0-0.02) | 0 | 0 | 0 (0-0.68) | 0 | 1389.32 (742.92-3624.01) | 0 | 0 (0-13.3) | 2.48 (0-3.25) | 0.4 (0.24-2.11) | 4.93 (0.72-18.79) | 2020.75 (1236.46-3817.63) | 13.63 (11.71-66.87) |
| | Mtb | 136.05 (0-210.11) | 0 (0-26.25) | 0.1 (0-0.27) | 1.47 (0.38-1.74) | 0 | 0.38 (0-2.44) | 13.08 (7.77-32.17) | 167.86 (39.26-396.99) | 0 (0-8.95) | 7.25 (0-26.74) | 6263.56 (1150.61-44609.43) | 6404.59 (4860.79-12430.25) | 0 | 0 (0-0.35) | 0 | 0 (0-33.16) | 1.84 (0.85-2.56) | 3.84 (0-7.48) | 2101.29 (612.36-4763.36) | 0 (0-1.57) | 5.53 (0-15.1) | 1.98 (0-6.06) | 1.12 (0-3.06) | 65.26 (8.96-243.99) | 869.5 (647.14-1537.01) | 68.33 (16.58-202.83) |
| | HIV-1 Bal | 505.03 (243.26-1459.9) | 31.92 (3.3-110.44) | 1.52 (0-2.1) | 1.14 (0.62-1.46) | 0 | 0.21 (0-0.8) | 1.8 (0.96-4.78) | 1.07 (0-2.07) | 0 (0-3.85) | 0 (0-18.93) | 239.47 (130.94-322.69) | 3629.26 (1572.5-7861.8) | 0 | 0 (0-2.95) | 0 | 0 (0-1.64) | 0.6 (0-1.18) | 0 (0-2.69) | 1919.28 (440.48-3023.7) | 0 (0-1.66) | 0 (0-15.32) | 1.67 (0-5.33) | 0.54 (0.21-1.76) | 39.12 (1.07-58.41) | 1856.64 (1213.2-2606.88) | 14.11 (11.45-115.68) |
| | Bal + Mtb | 158.92 (61.3-216.8) | 0 (0-38.47) | 0 (0-0.1) | 0.93 (0.38-1.3) | 0 | 0.41 (0-1.76) | 11.52 (6.26-64.97) | 158.06 (35.22-464.25) | 0 (0-7.46) | 8.37 (0-34.45) | 85538.66 (1455.25-365609.44) | 8218.54 (5864.24-15724.75) | 0 | 0 | 0 | 0 (0-5.85) | 1.49 (0.88-2.79) | 4.39 (0-5.06) | 1935.71 (578.09-5590.25) | 0 | 6.53 (0-27.35) | 1.71 (0-3.89) | 0.94 (0.4-1.44) | 61.56 (16.97-1444.92) | 976.78 (663.59-1361.95) | 42.66 (25.01-408.63) |
| | HIV-1 VSV-G | 154.13 (33.45-1165.27) | 0 (0-76.83) | 0.27 (0-2.1) | 0.73 (0.51-1.64) | 0 (0-4.96) | 0.47 (0-0.79) | 2.82 (1.41-4.82) | 1.92 (0-3.29) | 0 (0-4.88) | 0 (0-22.66) | 439.0 (262.130-638.91) | 4305.38 (3734.7-8460.84) | 0 | 0 (0-1.18) | 0 | 0 (0-0.67) | 0.68 (0-1.08) | 0 (0-3.3) | 2649.64 (284.64-3010.82) | 0 | 0 (0-17.76) | 1.67 (0-3.91) | 0.63 (0.21-1.14) | 13.73 (0-93.07) | 1535.38 (815.43-3774.65) | 13.26 (8.21-143.24) |
| | VSV-G + Mtb | 92.2 (0-232.15) | 0 (0-36.45) | 0 | 1.46 (0.26-1.74) | 0 (0-5.85) | 1.01 (0-3.22) | 10.34 (8.89-62.92) | 126.38 (50.98-545.91) | 0 (0-5.8) | 0 (0-58.47) | 4994.78 (3988.97-135984) | 8269.78 (4824.17-17363.99) | 0 | 0 (0-0.58) | 0 | 0 (0-16.12) | 2.33 (1.51-2.79) | 3.53 (0-6.73) | 1749.21 (302.77-5149.17) | 0 (0-2.64) | 6.65 (0-30.41) | 1.98 (0-4.68) | 0.87 (0.36-4.27) | 30.24 (4.37-1709.64) | 756.89 (648.77-1397.81) | 44.56 (14.09-570.52) |
| Pericytes | Uninfected | 3178.22 (2824.99-3192.33) | 270.97 (230.14-355.07) | 5.94 (4.67-7.53) | 0.28 (0-0.55) | 0 | 0 | 0 (0-0.57) | 0 | 0 | 0 | 94.55 (18.89-256.67) | 1238.77 (832.6-2333.01) | 0 | 0 | 0 | 0 | 0 (0-0.94) | 0 | 1823.72 (767.84-2554.32) | 0 | 6.27 (1.35-15.59) | 0 | 0 (0-0.68) | 4.39 (0.05-6.5) | 551.06 (308.45-790.14) | 1.09 (0-4.63) |
| | Mtb | 2589.47 (2482.42-2661.7) | 205.42 (179.96-277.9) | 4.35 (1.52-5.05) | 0.24 (0-0.34) | 0 | 0.31 (0.18-0.48) | 0.24 (0.16-1.1) | 0 | 0 | 0.71 (0-23.87) | 982.51 (564.92-1977) | 2155.21 (1389.59-5072.29) | 0 | 0 | 0 | 0 | 13.22 (11.47-20.77) | 19.8 (12.79-20.8) | 1929.58 (300.8-3110.5) | 0 | 0 (0-5.88) | 0 | 0 (0-0.79) | 122.87 (43.29-175.15) | 421.53 (346.54-677.74) | 27.77 (19.27-56.19) |
| | HIV-1 Bal | 2799.88 (2686.63-3066.3) | 203.48 (191.69-251.11) | 4.89 (4.35-6.12) | 0 (0-0.3) | 0 | 0 | 0.12 (0-0.37) | 0 | 0 | 0 | 10.3 (9.26-186.04) | 1010.67 (751.25-1844) | 0 | 0 | 0 | 0 | 0 | 0 | 1494.14 (583.67-2154.04) | 0 | 7.05 (4.02-18.18) | 0 | 0 (0-0.4) | 0 (0-1.42) | 286.85 (252.94-446.02) | 0.49 (0-1.16) |
| | Bal + Mtb | 2641.79 (2308.58-2855.07) | 189.69 (162.79-218.72) | 3.66 (2.62-3.81) | 0.12 (0-0.55) | 0 | 0.48 (0.26-1.05) | 0.41 (0.36-1.52) | 0 | 0 | 9.94 (0-14.41) | 955.7 (815.01-2688.37) | 2386.06 (1305.25-6255.09) | 0 | 0 | 0 | 0 | 12.96 (12.83-23.42) | 21.34 (15.47-23.28) | 1706.93 (422.73-2026.59) | 0 | 1.35 (0-10.1) | 0 | 0.44 (0-0.94) | 109.35 (84.48-172.23) | 477.3 (324.29-664.76) | 37.72 (24.19-42.51) |
| | HIV-1 VSV-G | 2394.74 (2320.2-2626.19) | 219.1 (178.16-233.95) | 3.66 (3.15-5.58) | 0.16 (0-0.44) | 0 | 0 | 0.14 (0-0.66) | 0 | 0 | 0 | 8.21 (3.56-16.96) | 1996.42 (878.7-2367.42) | 0 | 0 | 0 | 0 | 0 | 0 | 1829.36 (429.24-1993.69) | 0 | 5.59 (0-13.86) | 0 | 0 (0-0.56) | 0 (0-0.32) | 414.37 (312.23-621.41) | 0 (0-1.11) |
| | VSV-G + Mtb | 2322.52 (1964.07-2610.31) | 152.29 (131.81-215.21) | 2.08 (0.27-4.89) | 0.26 (0.08-0.65) | 0 | 0.67 (0.53-0.9) | 0.81 (0.5-1.34) | 0 | 0 | 18.3 (5.92-25.99) | 2657.64 (859.91-4731.6) | 2601.74 (1486.96-6720.19) | 0 | 0 | 0 | 0 | 20.03 (14.24-34.46) | 24.36 (17.59-36.99) | 1442.07 (419.66-2171.2) | 0 | 3.23 (0-8.11) | 0 | 0.68 (0.32-1.12) | 207.7 (80.54-233.8) | 658.96 (445.67-831.6) | 40.56 (28.54-60.55) |
| Microglia | Uninfected | 65.67 (54.5-141.05) | 40.23 (29.33-47.18) | 3.94 (2.08-7.48) | 12.84 (2.37-19.35) | 0 | 0 | 0 | 0 | 0 | 0 | 362.76 (297.67-566.88) | 185.61 (175.95-211.78) | 0 | 0 | 0 | 0 | 0 (0-0.87) | 0 | 1179.83 (723.59-1741.01) | 0 | 0 (0-1.17) | 0 (0-0.17) | 0 (0-0.56) | 0.31 (0-1.53) | 631.61 (199.49-1169.77) | 1.28 (0-2.73) |
| | Mtb | 33.45 (16.6-81.28) | 103.86 (98.39-215.11) | 4.67 (2.19-5.94) | 5.42 (1.05-13.74) | 0 | 0.3 (0-1.16) | 0.17 (0-0.21) | 0 | 0 (0-3.28) | 0 | 1598.29 (1037.37-5038.85) | 1800.65 (835.44-1919.26) | 0 | 0 | 0 | 0 | 19.14 (13.2-45.01) | 12.86 (8.62-36.38) | 1236.95 (1097.84-2561.12) | 0 | 0.02 (0-3.04) | 0 (0-0.32) | 0 (0-1.76) | 0.9 (0.5-5.17) | 446.08 (146.82-873.37) | 12.25 (11.11-23.34) |
| | HIV-1 Bal | 67.09 (53.56-207.04) | 24.17 (19.28-27.28) | 2.69 (1.52-4.35) | 7.75 (3.43-14.13) | 0 | 0 | 0 | 0 | 0 (0-1.04) | 0 | 439.65 (346.92-698.17) | 209.12 (163.88-251.66) | 0 | 0 | 0 | 0 | 0.63 (0.37-1.71) | 0 | 1130.12 (692.8-1796.16) | 0 | 0 | 0 | 0 (0-0.94) | 0.42 (0-2.04) | 535.35 (258.47-1098.53) | 2.84 (0.27-4.53) |
| | Bal + Mtb | 26.63 (16.6-195.64) | 131.39 (92.74-225.02) | 2.66 (1.56-5.64) | 3.93 (0.76-15.37) | 0 | 0.47 (0-0.56) | 0 (0-0.42) | 0 | 0 | 0 | 1684.16 (1495.22-7203.22) | 1453.97 (999.81-2050.4) | 0 | 0 | 0 | 0 | 16.11 (10.49-44.21) | 6.28 (5.3-40.38) | 1631.8 (1145.16-2669.59) | 0 | 0 (0-1.35) | 0 (0-0.53) | 0.24 (0-1.41) | 1.11 (0.55-7.05) | 365.35 (125.2-859.02) | 10.56 (7.62-23.03) |
| | HIV-1 VSV-G | 61.3 (34.74-283.68) | 26.25 (15.83-38.6) | 2.69 (0.27-5.58) | 11.10 (0.05-26.4) | 0 | 0 | 0 | 0 | 0 | 0 | 735.55 (573.53-807.16) | 464.79 (213-673.57) | 0 | 0 | 0 | 0 | 0.51 (0.21-11.54) | 0 | 1446.92 (677.17-1612.87) | 0 | 0 | 0 | 0.1 (0-0.56) | 0.52 (0-3.23) | 533.15 (308.77-997.99) | 2.28 (1.47-8.71) |
| | VSV-G + Mtb | 207.68 (0-355.04) | 118.92 (81.23-154.19) | 3.66 (1.56-6.59) | 4.8 (2.9-29.31) | 0 | 0.85 (0-1.2) | 0.32 (0-0.53) | 0 | 0 (0-3.85) | 0 | 1648.12 (1573.15-6893.05) | 2722.99 (1282.57-3622.5) | 0 | 0 | 0 | 0 | 25.48 (18.9-50.68) | 10.72 (2.86-29.97) | 1055.14 (772.45-1558.59) | 0 | 0 (0-1.17) | 0 (0-0.18) | 0.32 (0-1.07) | 1.11 (0.32-13.08) | 445.36 (194.98-1264.34) | 21.72 (9.06-32.59) |
| BBB | Uninfected | 2210.94 (1785.2-3611.21) | 179.75 (125.15-217.48) | 9.16 (7.21-40.54) | 1.27 (0.95-1.53) | 0 | 0.07 (0-0.17) | 0.67 (0.12-1.34) | 0 (0-0.03) | 0 (0-4.97) | 0 (0-2.56) | 404.88 (164.59-889.3) | 1996.42 (964.75-3590.53) | 0 | 0 (0-0.01) | 0 | 0 | 0.3 (0-1.12) | 0 | 3239.51 (2807.15-5296.08) | 0 | 12.53 (9.3-18.34) | 0.59 (0-0.91) | 0.94 (0.71-1.18) | 3.77 (1.06-9.19) | 1399.33 (1062.53-1563.08) | 9.66 (2.22-13.97) |
| | Mtb | 1606.53 (1476.95-2789.18) | 186.72 (119.5-216.08) | 10.92 (6.8-43.67) | 1.37 (1.05-1.61) | 0 | 0.21 (0.16-0.34) | 2.39 (1.25-3.78) | 5.21 (3.69-12.96) | 0 (0-6.81) | 0 | 1921.78 (1034.63-3566.02) | 4137.56 (2562.99-5605.20) | 0 | 0 (0-0.65) | 0 | 0 | 1.86 (0.88-8.78) | 0 (0-6.63) | 4003.25 (3165.47-6152.16) | 0 | 20.07 (16.43-21.65) | 0.92 (0-2.03) | 1.28 (0.87-2.18) | 10.75 (2.85-25.73) | 1377.18 (883.04-1475.81) | 12.53 (4.5-17.76) |
| | HIV-1 Bal | 2211.38 (1670.04-2713.66) | 173.75 (119.02-226.72) | 6.97 (5.5-45.31) | 1.23 (1.05-1.58) | 0 | 0.11 (0.05-0.57) | 0.69 (0.34-1.15) | 0 (0-0.12) | 0 (0-4.11) | 0 | 317.67 (246.14-771.26) | 2280.69 (1682.98-3235.73) | 0 | 0 (0-1.07) | 0 | 0 | 0.69 (0.31-1.24) | 0 | 3561.02 (2850.91-5778.18) | 0 | 14.41 (7.72-18.24) | 0 (0-1.86) | 1.07 (0.5-1.9) | 2.51 (2.06-7.05) | 1313.79 (1022.85-1526.64) | 6.79 (2.91-7.84) |
| | Bal + Mtb | 1874.97 (1417.09-2611.99) | 185.72 (106.13-248.57) | 7.66 (4.83-48.53) | 0.95 (0.83-1.95) | 0 | 0.32 (0.14-0.44) | 2.72 (1.58-4.02) | 7.68 (5.27-15.87) | 0 (0-6.23) | 0 (0-6.68) | 1724.52 (563.03-3224.38) | 4476.68 (2262.59-6322.28) | 0 | 0 (0-0.75) | 0 | 0 | 2.02 (0.99-11.7) | 0.48 (0-8.94) | 4368.67 (3416.18-4431.74) | 0 | 26.26 (13.04-28.16) | 0.7 (0.16-1.91) | 2.18 (1.36-2.83) | 10.72 (4.32-40.27) | 1188.75 (744.71-1339.49) | 9.43 (4.14-16.04) |
| | HIV-1 VSV-G | 1735.48 (1224.89-2531.74) | 134.9 (81.17-214.04) | 7.66 (5.5-43.6) | 1.42 (0.96-1.61) | 0 | 0.24 (0.17-0.45) | 0.62 (0.33-1.08) | 0 (0-0.04) | 0 (0-3.76) | 0 | 374.61 (245.76-764.84) | 2243.92 (1798.02-2942.66) | 0 | 0 (0-0.95) | 0 | 0 | 1.22 (0.4-1.28) | 0 | 3588.97 (3164.57-5829.05) | 0 | 9.89 (5.51-18.9) | 0.04 (0-1.67) | 0.3 (0-1.25) | 4.48 (1.25-9.77) | 1089.83 (839.35-1492.59) | 7.02 (3.37-9.41) |
| | VSV-G + Mtb | 1785.65 (1353.44-2559.03) | 160.64 (94.16-225.13) | 6.11 (3.19-46.05) | 1.12 (0.79-1.61) | 0 (0-1.41) | 0.23 (0.19-0.33) | 3.09 (1.44-4.39) | 9.77 (4.81-16) | 3.52 (0-8.4) | 0 (0-8.7) | 2125.96 (552.02-4415.93) | 5322.68 (3513.98-10338) | 0 | 0 | 0 | 0 | 2.88 (1.57-9.74) | 0 (0-6.18) | 4601.21 (3443.04-5449.21) | 0 | 25.53 (12.25-26.22) | 0.76 (0.06-2.96) | 1.76 (1.03-1.91) | 14.07 (6.9-42.44) | 1341.3 (646.67-1795.38) | 8.92 (4.13-18.63) |

**Supplementary Table 1. Modulation of soluble factors in CNS cells and BBB by *Mtb* and/or HIV-1 infection.**

| | | Astrocytes | | Pericytes | | Endothelial cells | | Microglia | | BBB | |
|---|---|---|---|---|---|---|---|---|---|---|---|
| | | **Mtb** | **HIV+Mtb** | **Mtb** | **HIV+Mtb** | **Mtb** | **HIV+Mtb** | **Mtb** | **HIV+Mtb** | **Mtb** | **HIV+Mtb** |
| Mitochondrial activity imbalance | | = | = | = | - (VSVG) | - | - (VSVG) | - | - (VSVG) | | |
| ROS release | | + | = | + | = | + | = | + | = | | |
| Extracellular glutamate | | + | = | + | + | = | = | + | = | + | = |
| MMP- | 2 | - | - (VSVG) | - | - (VSVG) | - | - (VSVG) | - | + (VSVG) | - | = |
| | 3 | = | = | - | - | - | = | + | + | = | = |
| Interleukins | 1β | = | = | = | = | + | = | = | = | + | + |
| | 6 | + | = | + | + (VSVG) | + | + (Bal) | + | = | + | + (VSVG) |
| | 8 | + | + (VSVG) | + | + | = | + | + | + (VSVG) | + | + (VSVG) |
| VEGF-A | | - | - | + | = | + | = | + | = | + | + |
| IP-10 | | + | + | + | + (VSVG) | + | = | = | = | + | + (VSVG) |
| MPC-1 | | - | = | - | + (VSVG) | - | - (VSVG) | - | - | = | = |
| IRE1 | XBP1 splicing | + | + | + | + | + | + | + | + (VSVG) | | |
| | RIDD activity | + | + (VSVG) | + | = | + | = | + | + (VSVG) | | |
| ATF6 | BiP expression | - | - | - | = | + | + | - | - (VSVG) | | |
| | uXBP1 expression | - | - (VSVG) | + | = | + | = | + | = | | |
| PERK | CHOP expression | + | = | + | + | + | + | + | = | | |
| Astrogliosis | C3 expression | + | + (Bal) | | | | | | | | |
| | S100A10 expression | = | + (VSVG) | | | | | | | | |
| Mtb entry | % | | + | | + | | = | | + | | |
| | MFI | | = | | = | | = | | = | | |
| Mtb growth | % | | + (VSVG) | | = | | = | | = | | |
| | MFI | | = | | + | | + | | = | | |
| BBB permeability | Integrity | | | | | | | | | - | - (Bal) |
| | CFU | | | | | | | | | | + |
| Cytotoxicity | Overall | + | + (VSVG) | + | + | + | = | + | = | | |
| | LT50 | | - (VSVG) | | - | | = | | = | | |
| Tight and adherens junctions | TJP1 | | | | | - | = | | | | |
| | F11R | | | | | - | - | | | | |
| | OCLN | | | | | - | = | | | | |
| | CLDN5 | | | | | - | - (VSVG) | | | | |
| | CDH5 | | | | | - | - | | | | |

**Supplementary Table 2. Summary of *Mtb* and/or HIV-1 effects on the BBB and CNS cells.**

| | Forward | Reverse |
|---|---|---|
| **Tight and adherens junctions** | | |
| TJP1 | 5′-AGAAGGATGTTTATCGTCGCATT-3′ | 5′-CCAAGAGCCCAGTTTTCCAT-3′ |
| F11R | 5′-CAAGTCGAGAGGAAACTGTTG-3′ | 5′-TCTGACTTCAGGTTCAGAAGAG-3′ |
| OCLN | 5′-GTCCAATATTTTGTGGGACAAGG-3′ | 5′-GGCACGTCCTGTGTGCCT -3′ |
| CLDN5 | 5′-GTGCTCTACCTGTTTTGCG-3′ | 5′-GACGGGTCGTAAAACTCG-3′ |
| CDH5 | 5′-CAGCCCAAAGTGTGTGAGAA-3′ | 5′-CGGTCAAACTGCCCATACTT-3′ |
| **Astrogliosis** | | |
| C3 | 5′-AAAAGGGGCGCAACAAGTTC-3′ | 5′-GATGCCTTCCGGGTTCTCAA-3′ |
| S100A10 | 5′-GGCTACTTAACAAAGGAGGACC-3′ | 5′-GAGGCCCGCAATTAGGGAAA-3′ |
| **Endoplasmic reticulum stress** | | |
| BLOS1 | 5′-CCCAATTTGCCAAGCAGACA-3′ | 5′-CATCCCCAATTTCCTTGAGTGC-3′ |
| sXBP1 | 5′-GCTGAGTCCGCAGCAGGT-3′ | 5′-CTGGGTCCAAGTTGTCCAGAAT-3′ |
| uXBP1 | 5′-CAGACTACGTGCACCTCTGC-3′ | |
| BiP | 5′-TCAGGCCAAGCCCAATACAG-3′ | 5′-TCCACGGTAGTGAGAGCCTT-3′ |
| CHOP | 5′-CAGAACCAGCAGAGGTCACA-3′ | 5′-AGCTGTGCCACTTTCCTTTC-3′ |
| **Housekeeping gene** | | |
| 18S | 5′-TAGAGGGACAAGTGGCGTTC-3′ | 5′-CGCTGAGCCAGTCAGTGT-3′ |

**Supplementary Table 3. List of primers used in the present study.**

**Supplementary Figure 1. Flow cytometry gating strategy.** Stained (eFluor$^{TM}$ fixable viability dye and FITC-conjugated anti-CD24 monoclonal antibody for cell viability and HIV-1, respectively), 4% PFA fixed and permeabilized cells were acquired on a LSRFortessa$^{TM}$ cell analyzer. A forward versus side scatter (FSC-A vs SSC-A) gate (called "Cells") is firstly used to identify the cell population and exclude debris. Onto the "Cells" gate, a forward scatter width (FSC-W) vs forward scatter height (FSC-H) gate (called "Singlets") is then used for multiplet exclusion. Onto the "Singlets" gate, a viability dye vs SSC-A gate (called "Alive") is then used to discern between living and dead cells. Finally, onto the "Alive" gate, a HIV-1 (FITC) vs *Mycobacterium tuberculosis* (RFP) quadrant gate is used to determine double negative (Mtb+HIV+), double positive (Mtb+HIV+) and single positive (Mtb+HIV- and Mtb-HIV+) cells. Compensation and fluorescence minus one (FMO) controls were used for every experiments.

**Supplementary Figure 2. Effect of *Mtb* and HIV-1 infection on endothelial cell tight cell, and adherens junction gene expression.** Endothelial cell gene expression of TJP1 (also known as ZO-1, panel A), F11R (encoding JAM-A, B), OCLN (Occludin, C), CLDN5 (Claudin-5, D) and CDH5 (encoding VE-Cadherin, E) was assessed by SYBR-Green Quantitative RT-PCR following infection with HIV-1 and/or *Mtb*. Results for 5 different experiments are presented as raw data normalized to the geometric mean of 18S ribosomal gene expression.

**Supplementary Figure 3. ROS assay positive control.** Uninfected astrocytes, pericytes, endothelial cells and microglia were treated with 50 µM Menadione for 4 hrs. before measure of $H_2O_2$. Data of 6 experiments are presented in raw data (i.e., RLU) in box and whiskers plots.

**Supplementary Figure 4. Effect of HIV-1 and/or *Mtb* infection(s) on ER stress.** sXBP1/uXBP1 (panels A-D), BLOC1S1 (E-H), CHOP (I-L), BiP (M-P), and uXBP1 (Q-T) transcripts were assessed by SYBR-Green Quantitative RT-PCR following infection(s) with *Mtb*, HIV-1 Bal and/or VSV-G pseudotyped HIV-1 of astrocytes (A, E, I, M, Q), hCMEC/d3 (B, F, J, N, R), HBVP (C, G, K, O, S), and HMC3 (D, H, L, P, T). Results for 5 to 6 different donors/passages are presented in raw data normalized to the geometric mean of 18S ribosomal gene expression in box and whiskers plots. Asterisks denote statistically significant data determined by ANOVA with Dunnett's correction for multiple comparisons (*$P < 0.05$, **$P < 0.01$, ***$P < 0.001$).

**Supplementary Table 1. Soluble factors released from CNS cells and the BBB model upon *Mtb* and/or HIV-1 infection.** The concentration of cytokines, chemokines, MMP and other soluble factors was measured after infection of CNS cells and BBB with *Mtb* and/or HIV-1 was determined by Luminex analysis as described in methods. The medians (IQR) of 7 (cells) or 5 (BBB) experiments are presented in raw data (i.e., pg/ml).

**Supplementary Table 2. Summary of *Mtb* and/or HIV-1 effects on the BBB and CNS cells.**
"*Mtb*" column: effect of *Mtb* infection compared to the uninfected control.

"HIV+*Mtb*" column: effect of HIV/*Mtb* co-infection compared to *Mtb* infection.

+ / - : increase / decrease

=: no effect or no increase

Bal / VSV-G: bystander effect / HIV-1 productive infection effect

**Supplementary Table 3. List of primers used in the study**
